# Mapping Functional Tumor Suppressor Networks in Esophageal Adenocarcinoma Using *In Vivo* CRISPR Screening and Perturb-sequencing

**DOI:** 10.64898/2026.07.19.739473

**Authors:** Ebtihal H Mustafa, Katherine Papastratos, KaMeng Wu, Niko Thio, Julia V Milne, Natasha Mitchell, Sasha Witts, Maree Pechlivanis, Bonnie Wong, Nicola Waddell, Wayne A Phillips, Nicholas J Clemons

**Affiliations:** Peter MacCallum Cancer Centre, Melbourne, VIC, 3000, Australia; Sir Peter MacCallum Department of Oncology, University of Melbourne, Parkville, VIC, 3010, Australia; QIMR Berghofer Medical Research Institute, Brisbane, QLD, 4006, Australia

## Abstract

Esophageal adenocarcinoma (EAC) is a genetically heterogeneous malignancy with few recurrent drivers, limiting effective targeted therapies. Although EAC arises from Barrett’s esophagus (BE), mechanisms driving progression from this premalignant state to invasive cancer remain unclear. We combined pooled CRISPR-Cas9 loss-of-function screening, *in vivo* tumorigenicity assays, and Perturb-seq profiling to define functional drivers of BE transformation. We identified 37 tumor suppressors whose loss promotes progression to EAC, defining a functional landscape of tumor initiation. Despite genetic diversity, these losses converged on four transcriptional programs involving metabolic reprogramming, cell cycle progression, RNA processing, and cellular motility. Furthermore, we identify loss of *NIPBL*, *TGFBR2*, and *RPL22* as key mediators of resistance to platinum- and taxane-based chemotherapy. Collectively, these findings provide a unifying framework for genomic heterogeneity in EAC, uncover underappreciated tumor suppressor pathways, and establish a resource to guide mechanistic and translational studies aimed at improving treatment strategies in this aggressive cancer.

## INTRODUCTION

Esophageal adenocarcinoma (EAC) represents a particularly intractable malignancy, characterized by frequent late diagnosis, limited therapeutic efficacy and consequently, poor clinical outcomes (Holmberg and Lagergren 2023, Li, Hoefnagel et al. 2023, Vissapragada, Bulamu et al. 2025). Progress toward developing molecular targeted treatments for EAC has been remarkably lacking. This stagnation stems largely from poor understanding of the fundamental mechanisms that underpin EAC tumorigenesis. Therefore, the comprehensive characterization of these core driver pathways is not merely advantageous but is, in fact, a required prerequisite for advancing new therapeutic interventions.

EAC arises from a premalignant condition known as Barrett’s esophagus (BE) (Song, Guha et al. 2007, Hvid-Jensen, Pedersen et al. 2011, Killcoyne and Fitzgerald 2021), which affects around 2% of the general population and develops as a result of chronic reflux disease (Song, Guha et al. 2007, Hvid-Jensen, Pedersen et al. 2011, Killcoyne and Fitzgerald 2021). BE is marked by the replacement of normal esophageal epithelium with a columnar, intestinal-like epithelium. Over time, this metaplastic tissue can progress to high-grade dysplasia (HGD) and eventually develops into invasive malignant lesions. Although this progression to EAC is well-recognized, our understanding of the molecular pathways driving this process remains incomplete.

Our limited, current understanding is largely inferred from whole-genome and exome sequencing studies of EAC, which have identified likely and potential driver genes (Dulak, Stojanov et al. 2013, Murugaesu, Wilson et al. 2015, Ross-Innes, Becq et al. 2015, Frankell, Jammula et al. 2019). These studies have highlighted: 1) an exceptionally high mutation rate, even in non-dysplastic BE, exceeding that observed in many other common cancers; 2) the identification of at least 76 putative driver genes in EAC, with an average of 4 to 5 potential driver events per tumor; 3) a notable lack of frequently recurring driver mutations; and 4) frequent copy number aberrations (CNAs).

Despite these insights, the molecular programs that precede transformation are largely obscured in established cancers. Thus, while this wealth of genomic knowledge holds promise for identifying novel therapeutic opportunities, there is a lack of functional studies to uncover the molecular pathways essential to EAC initiation and progression. Moreover, the data suggest that most EAC cases are driven by multiple molecular pathways, characterized by stochastic and heterogeneous low-frequency driver events (Frankell, Jammula et al. 2019). This complexity highlights the urgent need for innovative, high-throughput strategies to functionally dissect mechanisms of EAC tumorigenesis.

Traditional approaches for studying tumor-type-specific driver genes, such as genetically engineered mouse models, are neither high-throughput nor suitable for studying EAC tumorigenesis due to the lack of mouse models that accurately mimic Barrett’s metaplasia, the requisite starting point for this disease. We propose that human cell models derived from Barrett’s epithelium could serve as a viable alternative and previously demonstrated that inactivation of SMAD4 was sufficient to promote malignant transformation of non-tumorigenic dysplastic Barrett’s esophagus cell lines xenografted in immunocompromised mice (Gotovac, Kader et al. 2021). Herein, we combined patient derived dysplastic Barrett’s esophagus cell models with loss-of-function CRISPR-Cas9 screens and single-cell RNA sequencing (Perturb-seq) to enable high-throughput functional characterization of EAC driver genes. We reveal multiple tumor suppressor genes whose loss is sufficient to drive tumor initiation in dysplastic BE cells, including both established and previously underappreciated drivers. Despite the profound genomic heterogeneity of EAC, we show that diverse tumor suppressor alterations converge on conserved transcriptional states. These shared programs reveal common biological dependencies underlying distinct genetic lesions and identify vulnerabilities that extend beyond individual driver mutations. Furthermore, the association of specific tumor suppressor deficiencies with chemotherapy resistance highlights potential opportunities for patient stratification and therapeutic intervention. Together, these results suggest that targeting convergent downstream cellular states may provide a more effective strategy for overcoming the molecular complexity of EAC than focusing on individual genetic alterations.

## RESULTS

### Pooled CRISPR-Cas9 loss-of-function screen to uncover drivers of tumorigenicity in EAC

From published EAC genomic studies (Dulak, Stojanov et al. 2013, Murugaesu, Wilson et al. 2015, Ross-Innes, Becq et al. 2015, Frankell, Jammula et al. 2019), we compiled a list of 56 candidate tumor suppressor genes, whose inactivation through truncating mutations, deleterious missense variants, in-frame deletions, copy number loss or DNA methylation may contribute to tumor initiation. Although loss of function of these genes is predicted to drive EAC tumorigenesis, direct evidence confirming their functional roles remains limited. To investigate this, we performed a pooled CRISPR-Cas9 loss-of-function screen targeting these candidates in BE derived cell lines **(Fig. 1a)**. CPB and CPD high-grade dysplasia (HGD) Barrett’s cell lines, which harbor TP53 mutations but remain non-tumorigenic, were engineered to stably express Cas9-mCherry. Single-cell clones with high editing efficiency were isolated and designated Cas9-CPB and Cas9-CPD. Cas9 activity was validated using an EGFP reporter assay where sgRNA mediated disruption of EGFP caused loss of fluorescence.

**Figure 1.**
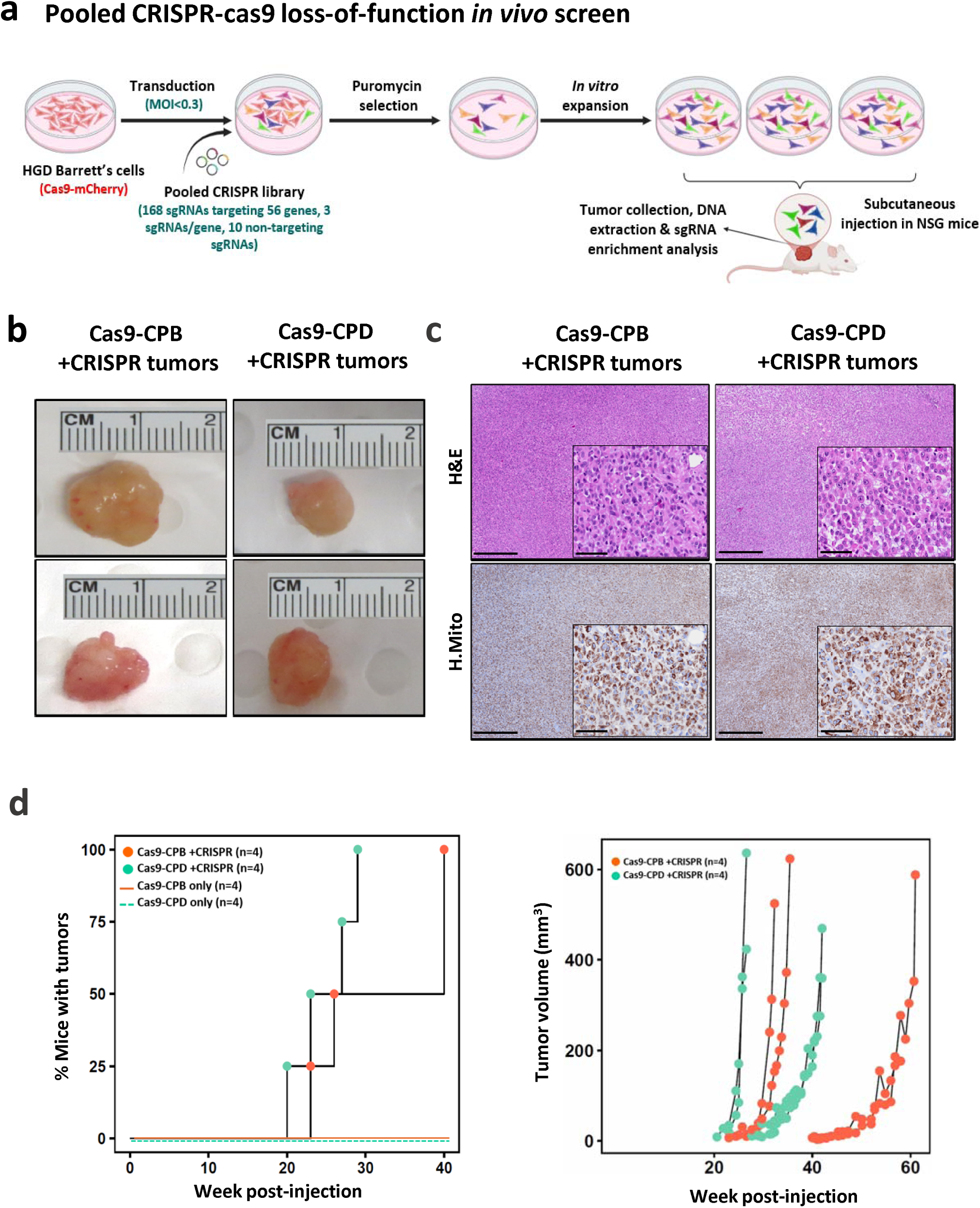
Pooled CRISPR-mediated knockout of putative EAC tumour suppressor genes promotes tumor formation from BE cells xenografted in NSG mice. **(a)** Schematic representation of the CRISPR loss-of-function screen in HGD Barrett’s esophagus cell lines, designed to identify tumor suppressor genes whose loss promotes transformation. **(b)** Representative images of tumours formed in NSG mice following transplantation of Barrett’s cells with the pooled CRISPR sgRNA library. **(c)** H&E staining and IHC for a human-specific mitochondrial protein (H.Mito). Scale bars, 100 µm (main) and 20 µm (insets). **(d)** Kaplan-Meier curve showing the proportion of mice developing tumours over time (left), alongside growth curves of individual tumours (right). Blue and red lines in the Kaplan-Meier graph indicate that mice transplanted with Cas9-CPB and Cas9-CPD cells, not subjected to CRISPR library transduction, did not develop tumours.

Flow cytometry confirmed Cas9 efficiencies greater than 95% in both clones **(Supplementary Fig. S1a)**, supporting their suitability for *in vivo* tumorigenicity experiments. These clones were transduced with a pooled CRISPR-Cas9 library comprising 178 dual sgRNA expression plasmids, each expressing two distinct sgRNAs targeting the same gene with three sgRNA pairs per gene, along with 10 non-targeting control plasmids. Tumours developed in all mice injected with Cas9-CPB and Cas9-CPD cells transduced with the pooled sgRNA library **(Fig. 1b-d)**, with a median latency of ∼20 weeks **(Fig. 1d)**. In contrast, and consistent with our previous findings (Gotovac, Kader et al. 2021), mice injected with Cas9-only CPB or CPD cells did not develop tumours **(Fig. 1d, Supplementary Fig. S1b,c)**, demonstrating that tumorigenesis was specifically driven by gene perturbations introduced through the CRISPR library.

To identify drivers of tumor formation, we profiled sgRNA representation across all tumours **(Fig. 2, Supplementary Fig. S2)**. On average, 52 sgRNAs targeting 34 genes were enriched per tumor. Shared sgRNA signatures demonstrated strong reproducibility **(Fig. 2a-c, Supplementary Fig. S2a-c)**, with 42 and 46 sgRNAs detected in three or more tumours from Cas9-CPB and Cas9-CPD cells targeting 37 and 31 distinct genes, respectively. Approximately 55% of these genes were targeted by multiple independent sgRNAs in at least one tumor **(Fig. 2d, Supplementary Fig. S2d)**. We found strong enrichment of sgRNAs targeting *PTEN* across all tumours **(Fig. 2b,c, Supplementary Fig. S2b,c)**, consistent with the well-established tumorigenic role of *PTEN* loss in multiple cancer types (Chalhoub and Baker 2009). Among the recurrently targeted genes identified across tumours derived from both BE cell lines and represented by multiple sgRNAs were *STK11*, a serine/threonine kinase that orchestrates cellular energy metabolism (Faubert, Vincent et al. 2014); the MAPK signalling mediators *MAP2K7* and *MAP3K1*; the Eph receptor family member *EPHA3* ; *DNAH7*, a force-generating axonemal dynein critical for ciliary function (Zhang, O’Neal et al. 2002), and *GPATCH8*, a putative regulator of RNA splicing (Karika, Cipakova et al. 2026).

**Figure 2.**
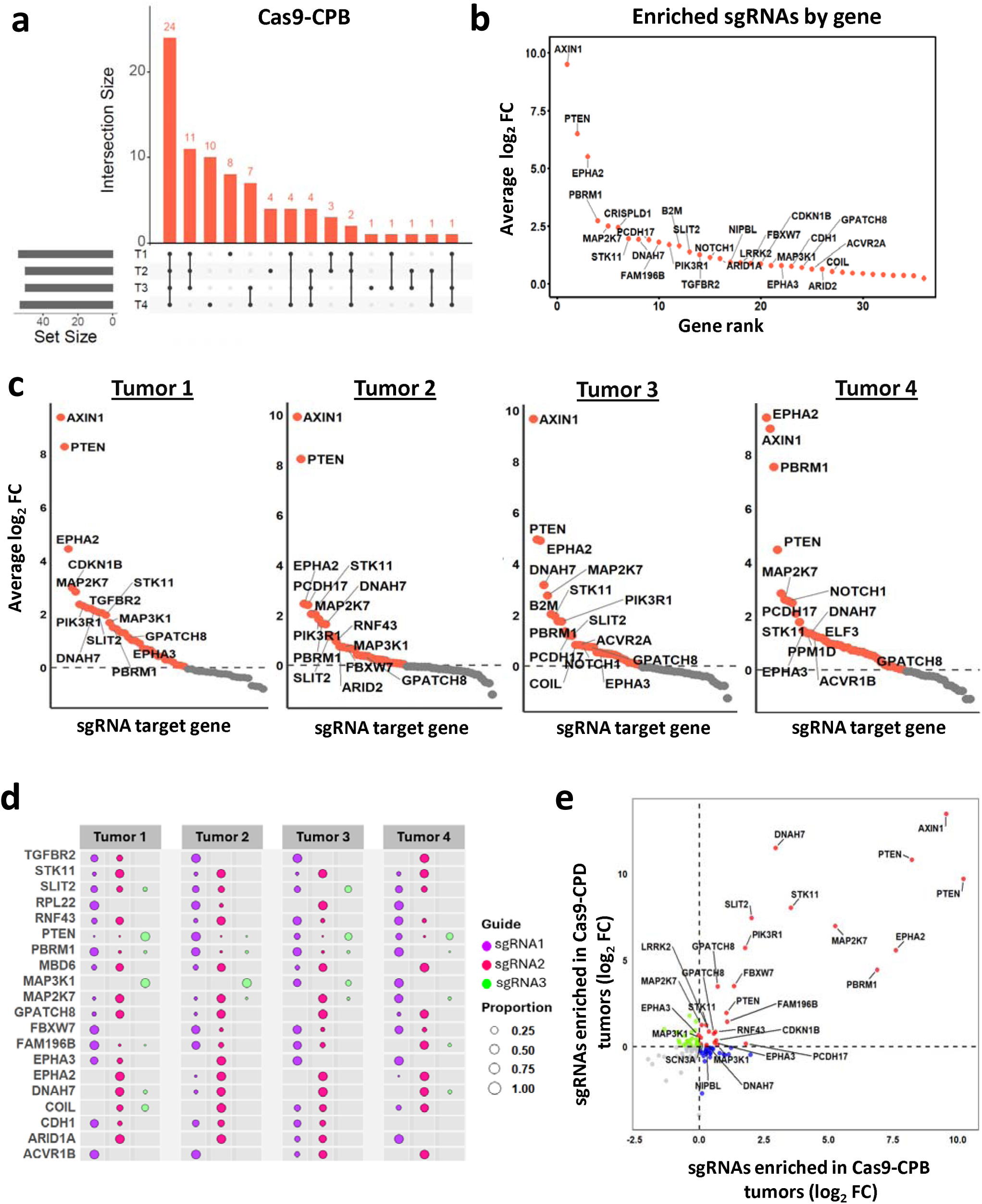
Identification of tumor suppressors whose loss drives BE cell transformation. **(a)** UpSet plot showing the number of enriched sgRNAs shared across four Cas9-CPB tumours. **(b)** Average log_2_ fold change (FC) of tumor-enriched sgRNAs in Cas9-CPB tumours, grouped by gene and relative to baseline (T0) cells. Displayed genes were identified in tumours from at least three mice. **(c)** MAGeCK plots showing log_2_ FC of enriched sgRNAs grouped by gene in four independent Cas9-CPB tumours. Top-enriched genes are labelled. **(d)** Enrichment of independent sgRNA clones across four tumours, with each gene targeted by three pairs of sgRNA. Dot size reflects the proportion of total reads per gene. **(e)** Two-dimensional enrichment (2D MA) plot comparing sgRNA enrichment between Cas9-CPB and Cas9-CPD tumours. Guides enriched in both cell lines cluster in the upper right quadrant (red). Blue dots indicate sgRNAs enriched only in Cas9-CPB tumours, green dots indicate those enriched only in Cas9-CPD tumours, and sgRNAs depleted in both models cluster in the lower left quadrant (grey). Repeated gene labels reflect independent enrichment by multiple independent pairs of sgRNAs.

### Profiling transcriptional effects of EAC driver genetic perturbations reveals convergent driver phenotypes

To identify the transcriptomic changes associated with perturbation of candidate driver genes in the BE cells and delineate the molecular signalling networks they influence, we applied a Perturb-seq technology (Dixit, Parnas et al. 2016), that integrates pooled CRISPR-Cas9-mediated knockouts with single-cell RNA sequencing **(Fig. 3a)**. This approach allowed us to examine the molecular effects of each perturbation in a high-throughput manner and was designed to functionally interrogate candidate genes emerging from our *in vivo* screen as drivers of EAC tumor initiation. However, we did not restrict our analysis to only those perturbations enriched in the *in vivo* screen but included all genes represented in the CRISPR library. While some of these genes were not identified in the *in vivo* screen, potentially because they do not directly promote tumor initiation on their own, characterizing their transcriptional effects can still provide important insights into their roles in tumor progression. This broader approach enabled a more comprehensive exploration of transcriptional programs altered by diverse EAC driver mutation events and facilitated the identification of distinct functional phenotypes through integrative clustering of perturbation-induced transcriptional profiles. The same CRISPR library used in our *in vivo* CRISPR screen was also used for Perturb-seq, where each sgRNA contained a unique capture sequence (CS) compatible with the 10x Genomics 31 Single Cell RNA-seq platform **(Fig. 3a)**, enabling accurate identification of individual perturbations at single-cell resolution.

**Figure 3.**
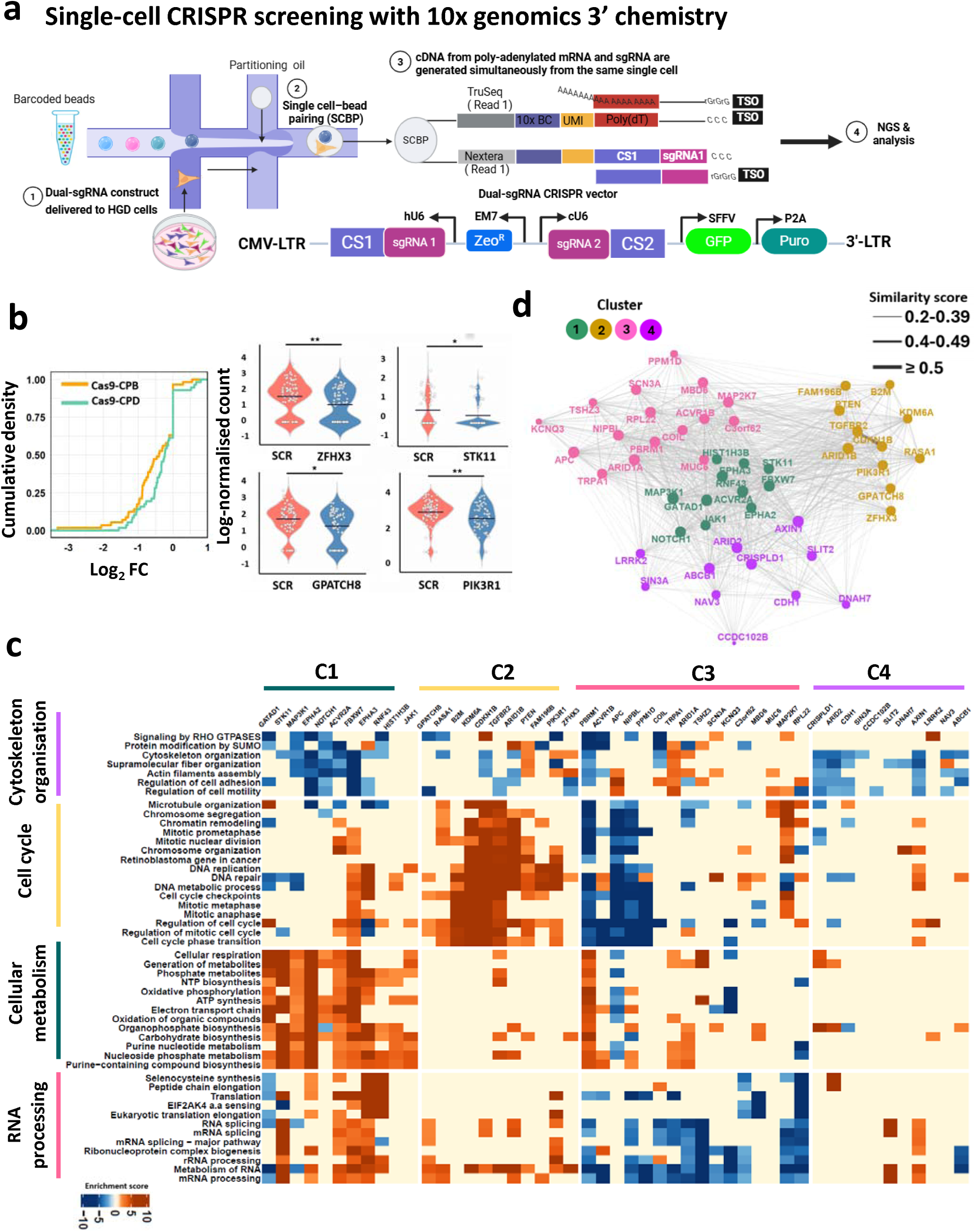
High-throughput Perturb-seq profiling of CRISPR-induced transcriptional changes. **(a)** Schematic of the Perturb-seq experiment. HGD Barrett’s cells were transduced with a pooled dual-sgRNA CRISPR library containing capture sequences (CS) compatible with 10X Genomics. Single cells were encapsulated with barcoded beads to capture both sgRNAs and polyA-mRNA, followed by library preparation and sequencing. **(b)** Cumulative density plots (left) showing the distribution of log_2_ FC for genes targeted by the sgRNA library and representative violin plots (right) showing log-normalized counts of selected targeted genes, reflecting reduced expression levels upon CRISPR-mediated knockout. *P-value* * < 0.05, ** < 0.001. **(c)** Heatmap showing clustering of perturbations based on the top enriched pathways. The Enrichment Score corresponds to log10(FDR-values), with positive values indicating enrichment in upregulated genes and negative values indicating enrichment in downregulated genes. **(d)** Network depicts similarity between perturbations based on shared enriched pathways, highlighting groups with related transcriptional programs. Nodes represent perturbations and are coloured according to Louvain clusters. Edges represent pairwise Jaccard similarity, with thickness proportional to the degree of similarity between the two genes.

After applying quality control filtering (>1,000 detected genes, >10,000 unique molecular identifiers, <5% mitochondrial gene content, and a dual-sgRNA construct targeting a single gene, **Supplementary Fig. S3a, b**), we retained 3,553 Cas9-CPB cells and 3,719 Cas9-CPD cells for downstream analysis. The cumulative density plot **(Fig. 3b)** shows the efficiency of CRISPR perturbations, where a leftward shift toward negative log2 fold change indicates that a large proportion of sgRNAs resulted in reduced expression of their target genes compared to scrambled controls. There was considerable variation in the number of cells per gene perturbation, ranging from 2 to 176 cells, with a median of 45 cells **(Supplementary Fig. S3c)**. Some perturbations were highly represented, which may reflect differences in cell viability following gene knockout, or may be due to technical factors such as variability in library pooling. To investigate how CRISPR perturbations reshape the transcriptional landscape, we performed a pathway enrichment analysis on differentially expressed genes **(Fig. 3c, Supplementary Fig. S3d)**. Clustering of the top enriched pathways demonstrated a clear convergence of the tumour suppressors into four distinct groups based on their shared transcriptional programs **(Fig. 3c)**; each fundamentally aligned with an established cancer hallmark. Broadly, the four transcriptional programs cover: (i) cell cycle division, regulation and progression; (ii) cell metabolism; (iii) RNA processing, translation, and protein stability; and (iv) cell adhesion and motility.

To confirm the reliability of our analysis pipeline, we examined the altered transcriptional pathways of representative examples of perturbed genes with well-known biological roles (**Supplementary Fig S4)**. Loss of *APC*, a negative regulator of WNT signalling, resulted in the upregulation of WNT pathway-associated genes, which in turn activated downstream processes like epithelial-mesenchymal transition and cell morphogenesis. Similarly, knockout of *RPL22*, a member of the ribosomal protein L22 family, led to a strong downregulation of pathways related to RNA splicing, RNA metabolic regulation, 60S ribosomal subunit assembly, and protein stability. Finally, knockout of *RASA1*, a negative regulator of RAS/MAPK and PI3K/AKT signalling, resulted in increased expression of genes involved in these pathways. Collectively, these representative examples confirm that our analysis pipeline reliably captures the transcriptional signatures associated with individual gene perturbations.

To complement the high-level summary provided by the clustering of transcriptional phenotypes **(Fig. 3c)** and fully capture the complex relationships between perturbations, we constructed a comprehensive network analysis using all significantly enriched pathways for each perturbation **(Fig. 3d, Supplementary Table 2)**. This approach highlights not only gene-level similarities but also reveals broader clusters of perturbations that converge on shared transcriptional programs. Cluster 1 is enriched for perturbations associated with metabolic reprogramming, driving expression of genes involved in aerobic respiration and respiratory electron transport, organophosphate biosynthetic process, metabolism of amino acids and derivatives, carboxylic acid metabolic process, metabolism of lipids, and others **(Fig. 3c, Supplementary Table 2)**. Well-established metabolic regulators such as *STK11* and *FBXW7* (Shimizu, Nihira et al. 2018, Minor, Couser et al. 2025), were identified within Cluster 1, together with *MAP3K1*, *EPHA3*, *EPHA2*, *NOTCH1*, *HIST1H3B*, *GATAD1*, and *ACVR2A*, which also shared effects on metabolic rewiring. Perturbation of Cluster 1 driver genes was also associated with altered expression of genes involved in actin cytoskeleton organisation, and cell polarity. Cluster 2 predominantly captured gene perturbations affecting cell cycle regulation. Loss of established cell cycle regulators, including *PTEN*, *CDKN1B*, and *TGFBR2* (Rojas, Padidam et al. 2009, Massacci, Perfetto et al. 2023, Khasarah, Nabilsi et al. 2025), led to pronounced upregulation of genes controlling sister chromatid segregation, mitotic progression, and cell cycle checkpoints. This transcriptional phenotype was recapitulated by inactivation of other genes, such as *PIK3R1*, *KDM6A*, *B2M*, *ARID1B*, and *RASA1*, which also influenced DNA processing and repair pathways. Interestingly, this cluster included a less well-characterised gene, *FAM196B*. Cluster 3 comprised genes with pronounced effects on RNA metabolism and processing, RNA splicing, eukaryotic translation, mitochondrial translation, protein modifications, and ribonucleoprotein synthesis **(Fig. 3c, Supplementary Table 2)**. This group included *RPL22*, *TSHZ3*, *C3orf62*, *TRPA1*, *KCNQ3*, and *ARID1A* genes. Chromatin re-modelers, including *ACVR1B*, *PBRM1*, *NIPBL*, *APC*, and *PPM1D*, also regulate these pathways and additionally influence cell cycle progression, DNA replication, and metabolic homeostasis. Finally, Cluster 4 was predominantly characterized by perturbations in actin cytoskeleton organization, cell adhesion, and cell motility. This cluster included *CDH1*, *NAV3*, *AXIN1*, *DNAH7*, and *ARID2* genes, most of which have previously been implicated in regulating cytoskeletal dynamics, migration, epithelial-mesenchymal transition, and invasive cell behaviour (Stringham and Schmidt 2009, Svitkina 2018, Qin, Wu et al. 2020, Moreno, Monterde et al. 2021).

To place the identified driver gene transcriptional clusters within a clinical context, a cross-cohort analysis was performed using the ICGC and TCGA (Firehose Legacy) EAC clinical datasets (n = 200, and 89, respectively), all of which are supported by comprehensive mutational and clinical metadata. The analysis focused on curated loss-of-function (LoF) alterations, including missense, nonsense, frameshift, and splice-site mutations, as well as deep deletions, to evaluate patterns of mutational exclusivity across the clusters. Cluster 1 genes, which showed mutation frequencies of 29% (ICGC), and 30.3% (TCGA) **(Supplementary Fig. S5a, S6a)**, demonstrated a high degree of mutual exclusivity, with independent LoF events occurring in more than 80% of the patients. This strong exclusionary pattern suggests potential tumor suppressor functional redundancy among the genes within this cluster. Similar results were observed in Cluster 2 (prevalence: 17.5% ICGC, and 40.4% TCGA, **Supplementary Fig. S5a, S6a**), which maintained exclusivity rates ranging from 70% to 88.6%. Conversely, Cluster 3 genes (prevalence: 55% ICGC, and 64% TCGA) showed lower exclusivity (60-67%), which could indicate a tendency for concurrent inactivation of multiple tumor suppressor genes within this cluster. Finally, Cluster 4 (prevalence: 39% ICGC, and 50.6% TCGA) showed substantial inter-cohort variability in exclusivity (71 % in ICGC and 49% in TCGA), highlighting divergent patterns of mutational co-occurrence across the analysed clinical populations. Despite these distinct genomic patterns, survival analysis demonstrated no significant difference in overall survival among the four transcriptional clusters, suggesting that their underlying genomic and transcriptional heterogeneity does not confer a differential prognostic outcome in EAC cohorts **(Supplementary Fig. S5b, S6b)**.

#### Loss of *PTEN, EPHA3, GPATCH8*, and *STK11* drives EAC initiation

To validate candidate drivers identified in our pooled CRISPR-Cas9 screen, we conducted single-gene loss-of-function experiments. *PTEN*, *EPHA3*, *GPATCH8*, and STK11 were selected based on their differing levels of enrichment across tumours, spanning high (*PTEN*), moderate (*STK11*), and lower enrichment (*EPHA3* and *GPATCH8*), as well as representation by multiple independent sgRNAs. Loss of single candidate tumor suppressor genes markedly enhanced tumor formation **(Fig. 4a, b)**. All mice injected with BE cells harbouring *PTEN*, *GPATCH8* or *EPHA3* knockouts developed tumours, with tumor onset occurring within a substantially reduced latency window (∼14-23 weeks post-injection), consistent with the median latency observed in the pooled CRISPR screen and supporting their role as robust EAC tumor drivers. Similarly, *STK11* knockout also resulted in tumor formation in injected mice; however, tumor development occurred with comparatively longer latency than the other gene knockouts **(Fig. 4b)**. PTEN was included in the *in vivo* validation experiments as a positive control, demonstrating that loss of a canonical tumor suppressor is sufficient to induce tumorigenicity. Consistent with its established biological function, transcriptomic analysis, demonstrated that *PTEN* loss induces tumorigenic transcriptional programs associated with activation of the PI3K/AKT signalling pathway **(Supplementary Fig. S7a)**.

**Figure 4.**
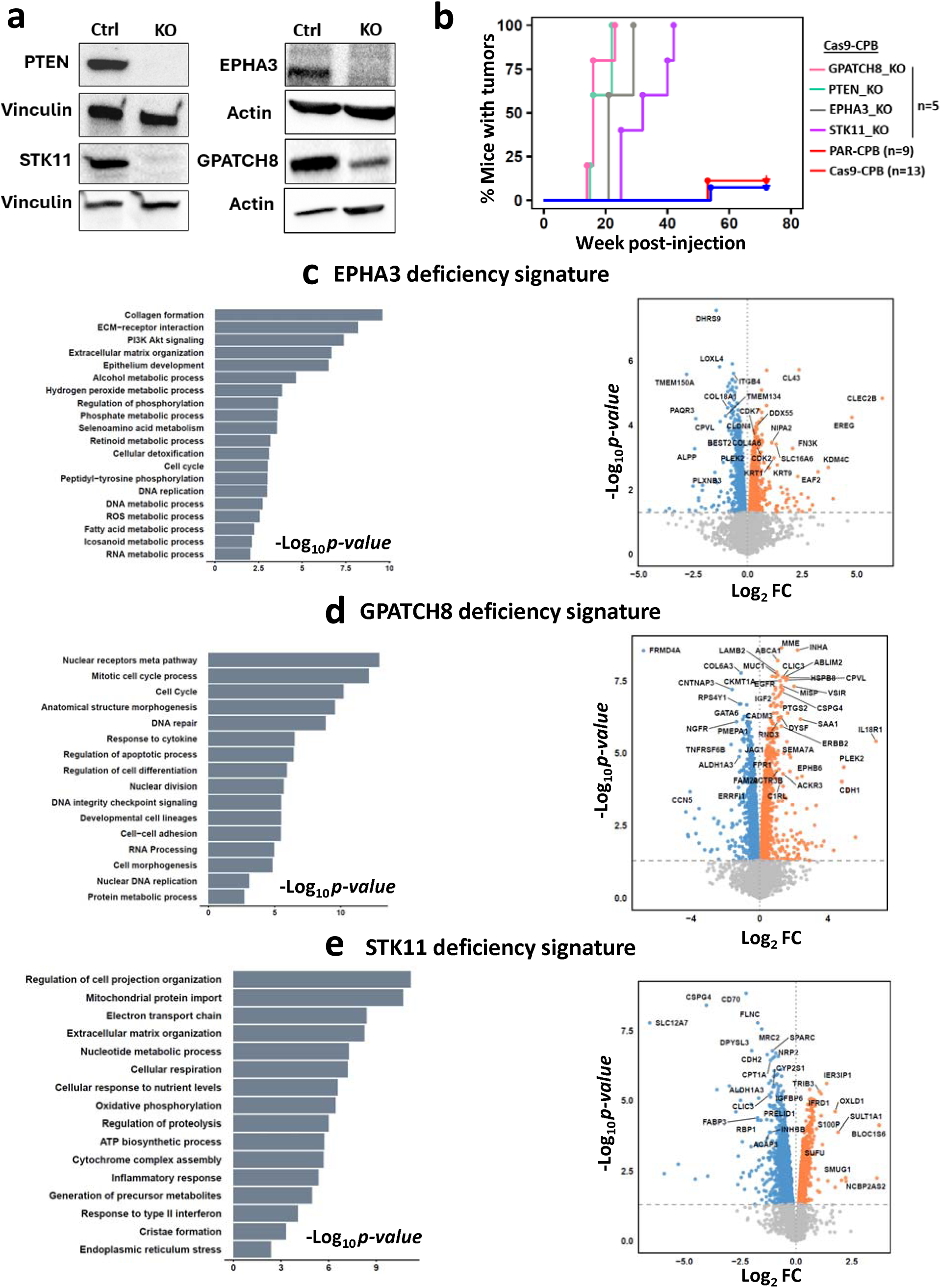
Loss of PTEN, EPHA3, GPATCH8, and STK11 drives tumor formation from BE cells. **(a)** Western blots for PTEN, EPHA3, GPATCH8, and STK11 protein in Cas9-CPB cells following CRISPR-mediated knockout. **(b)** Kaplan-Meier curve illustrating the proportion of mice developing tumours over time following knockout of *PTEN*, *EPHA3*, *GPATCH8*, and *STK11* in Cas9-CPB cells, compared with parental CP-B and Cas9-CPB control groups. **(c-e)** Bar and volcano plots showing pathway enrichment analysis of differentially expressed proteins from Cas9-CPB cells with *EPHA3, GPATCH8, and STK11* knockout compared with Cas9-CPB control cells.

To define the molecular programs driven by EPHA3, GPATCH8, and STK11 knockout and assess whether Perturb-seq phenotypes were reflected at the protein level, we expanded our analysis to include quantitative proteomic profiling **(Fig. 4c, d, e, Supplementary Table 3)**. Protein-level analysis of CP-B Barrett’s cells following *EPHA3* knockout demonstrated strong concordance with the Perturb-seq transcriptional signatures **(Fig. 3c, Fig. 4c)**. Pathway enrichment analysis of differentially expressed proteins showed enrichment of cellular metabolic processes, including phosphate, seleno-amino acid and nucleic acid metabolism, together with alterations in extracellular matrix organization, collagen formation, and focal adhesion pathways **(Fig. 4c)**. Proteomic profiling of GPATCH8-deficient Barrett’s CP-B cells extended the findings from the single-cell RNA sequencing analysis, revealing broad dysregulation of biological programs involved in nuclear receptor signalling, cell-cycle regulation, DNA repair, RNA processing, and cell morphogenesis **(Fig. 4d)**. Knockout of *STK11* in CP-B Barrett’s cells, as captured by both Perturb-seq and proteomic analyses **(Fig. 3c, Fig. 4e)**, resulted in coordinated alterations in metabolic, stress-response, and inflammatory pathways. Collectively, our findings establish *PTEN*, *EPHA3*, *GPATCH8*, and *STK11* as tumor suppressors that promote EAC development when lost and provide additional support for the molecular phenotypes identified by Perturb-seq.

### Pooled CRISPR-Cas9 screening highlights drivers of innate resistance to Carboplatin and Paclitaxel

To further characterize the functional consequences of tumor suppressor gene loss in EAC, we investigated whether any of these alterations were associated with resistance to the standard-of-care chemotherapies carboplatin and paclitaxel (Van Hagen, Hulshof et al. 2012). Carboplatin acts by forming DNA crosslinks that interfere with replication and transcription, ultimately leading to cell death (Sousa, Wlodarczyk et al. 2014), while paclitaxel is a microtubule-stabilizing agent that disrupts mitotic spindle function and induces mitotic arrest (Foland, Dentler et al. 2005). The same cell line models transduced with our pooled CRISPR knockout library, were challenged with cytotoxic drug concentrations **(Fig. 5a)**, chosen based on prior cell viability experiments and confirmed to kill over 70% of the parental cell population **(Supplementary Fig. S7b, c)**, thus imposing strong selective pressure. Comparative analysis of chemotherapy-treated versus vehicle-treated populations identified tumor suppressors associated with resistance **(Supplementary Fig. S7d, e)**. Focusing on the resistance-conferring gene knockouts shared by both cell lines, we specifically identified six genes whose loss mediated dual resistance to both paclitaxel and carboplatin **(Fig. 5b, red dots)**. This set comprises: the chromatin re-modelling factors *NIPBL* and *PBRM1*; the WNT pathway negative regulators *AXIN1* and *RNF43*; *TGFBR2*; and *FAM196B*. We also identified tumor suppressors whose loss conferred resistance specifically to paclitaxel (*CRISPLD1*, *SLIT2*, *APC*, and *CDKN1B*) or carboplatin (*RPL22*, *ARID1A*, *MAP2K7*, and *ACVR1B*). We next assessed the clinical relevance of these chemoresistance-associated genetic alterations in EAC patient cohorts (ICGC and TCGA data). Approximately 46% of patients harboured a mutation in at least one of these genes, while 18% carried alterations in at least one of the six common tumour suppressors identified across both carboplatin and paclitaxel resistance screens **(Fig. 5c)**. To explore the clinical relevance of these findings, we analysed transcriptomic and survival data from 92 patients with EAC enrolled in the DOCTOR phase II clinical trial (ANZCTR - ACTRN12609000665235), in which all patients received docetaxel (paclitaxel analogue), cisplatin (carboplatin analogue), and 5-fluorouracil (DCF). Patients with low *NIPBL* expression, defined as expression below the median log2 counts per million (CPM), showed significantly poorer overall survival compared with patients with high *NIPBL* expression **(Fig. 5d)**. *NIPBL* was the top common hit identified in both the carboplatin and paclitaxel resistance screens. Similar associations with reduced overall survival were also observed in patients with low expression of *TGFBR2* and *RPL22* **(Fig. 5d)**. Together, these findings identify recurrent tumour suppressor losses that mediate resistance to standard chemotherapies in EAC and highlight loss of *NIPBL*, *TGFBR2*, and *RPL22* as potential biomarkers of resistance.

**Figure 5.**
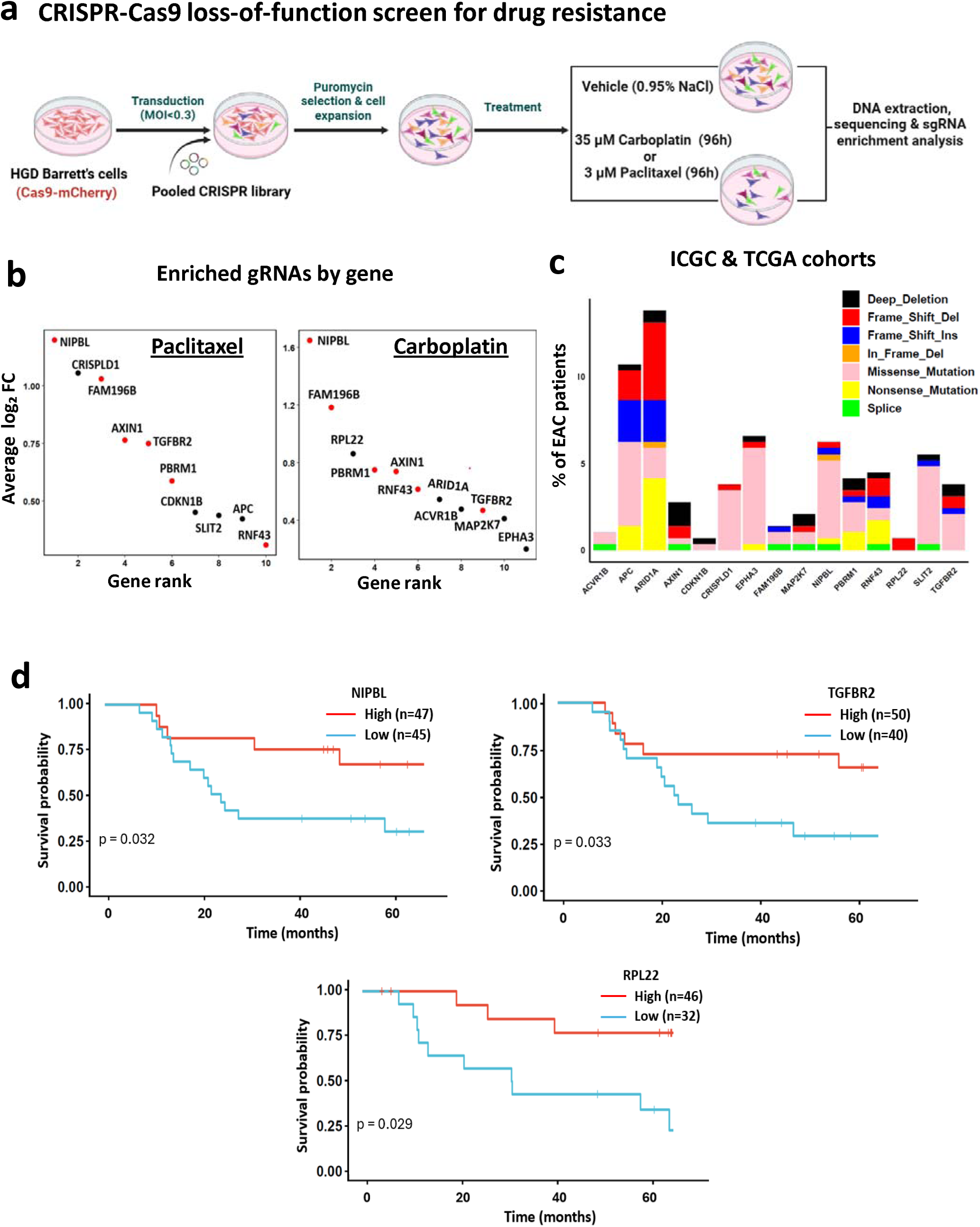
CRISPR-Cas9 screen reveals genetic drivers of intrinsic resistance to carboplatin and paclitaxel. **(a)** Schematic of the drug-screen experiment. CRISPR-engineered HGD Barrett’s cells were treated with vehicle (0.9% saline), 35 µM carboplatin, or 3 µM paclitaxel for 96 hours, followed by DNA extraction and sgRNA sequencing to identify enriched guides. **(b)** Average log_2_ FC of enriched sgRNAs following paclitaxel or carboplatin treatment, grouped by gene and compared with vehicle-treated cells. Red dots indicate genes whose loss confers resistance to both drugs. **(c)** Bar chart showing the percentage of EAC patients in the ICGC and TCGA cohorts with deleterious mutations or deep deletions in these resistance-associated genes. **(d)** Kaplan-Meier curves showing overall survival of EAC patients from the DOCTOR trial treated with DCF chemotherapy in relation to mRNA expression levels of *NIPBL*, *TGFBR2*, and *RPL22*. Low and high mRNA expression groups were defined using the median log2 CPM value as the cutoff, where low expression was ≤ median and high expression was > median.

## DISCUSSION

Our understanding of the molecular genetic events that drive progression from Barrett’s esophagus to invasive EAC, a highly lethal malignancy, has been limited by a lack of detailed functional based studies. In this study, we combined pooled CRISPR-Cas9 loss-of-function screening, *in vivo* tumorigenicity assays, Perturb-seq transcriptional profiling, and chemotherapy resistance screening to define functional tumor suppressors in human-derived models of EAC initiation. Together, our findings provide functional evidence to define tumor suppressor genes that actively drive EAC tumorigenesis. We show that perturbation of diverse tumor suppressor genes converges into shared transcriptional programs and establish key tumor suppressors that influence intrinsic resistance to standard chemotherapeutic agents.

A key finding emerging from the Perturb-seq analysis was the convergence of diverse EAC driver gene perturbations into four major transcriptional programs broadly associated with canonical cancer hallmarks: metabolic reprogramming, cell cycle progression, RNA processing and translation, and cell motility. Despite the genetic heterogeneity of EAC, these results suggest that many tumor suppressor alterations ultimately funnel into a limited number of functional cellular states. Such convergence may help explain how genetically diverse tumours acquire similar malignant phenotypes and may provide a framework for identifying shared therapeutically targetable biological dependencies.

Importantly, integration of our transcriptional clusters with clinical EAC datasets demonstrated that these functional gene groups exhibit distinct mutational profiles in patient tumours. Clusters associated with metabolic regulation and cell cycle control showed high degrees of mutual exclusivity, suggesting functional redundancy among genes converging on similar biological programs. In contrast, RNA-processing-associated perturbations displayed greater co-occurrence, potentially reflecting additive or cooperative effects on cellular fitness. Despite these differences, we did not observe significant survival differences between clusters in the ICGC and TCGA cohort. This may reflect the multifactorial nature of EAC progression and treatment response, or the possibility that distinct molecular routes ultimately converge on similarly aggressive disease phenotypes.

Importantly, although our transcriptional clustering analysis demonstrated that functionally related perturbations also show convergent mutational patterns in independent clinical EAC cohorts, direct validation of these transcriptional states in patient-derived samples during progression remains challenging. An ideal comparison would require matched dysplastic Barrett’s esophagus and EAC tissues collected longitudinally from the same patients, with paired whole-genome and transcriptomic sequencing data available across disease progression. Such datasets remain extremely limited. In our experimental system, we modelled this evolutionary process using isogenic Barrett’s-derived epithelial cells in which defined tumor suppressor alterations were introduced individually, thereby enabling direct attribution of transcriptional changes to specific genetic perturbations without the confounding effect of interpatient background genetic variability. This strategy effectively mimics the transition from premalignant Barrett’s epithelium toward malignancy under controlled conditions and allowed us to identify transcriptional programs associated with recurrent EAC-associated mutations. Although this approach cannot fully capture the complexity of tumor evolution or the influence of the tumor microenvironment in patients, the observation that these experimentally derived transcriptional clusters are also reflected within independent clinical cohorts strongly supports their biological relevance.

Our pooled *in vivo* CRISPR-Cas9 screen identified 37 tumor suppressor genes whose disruption promoted tumor formation in Barrett’s-derived epithelial cells. Among the genes consistently enriched in our screen were *PTEN*, *STK11*, *MAP2K7*, *MAP3K1*, *EPHA3*, *EPHA2*, *GPATCH8*, *AXIN1*, *PIK3R1*, and *PBRM1*. We selected a subset of candidates for single-gene validation experiments, including genes with well-established tumor-suppressive roles in other cancers such as *PTEN* and *STK11* (Carretero, Medina et al. 2004, Marinaccio, Suraneni et al. 2021, Khasarah, Nabilsi et al. 2025), a gene (EPHA3) with a context-dependent or dual role in cancer (Lv, Wang et al. 2018, Chen, Lu et al. 2019), and a gene with a less characterized function, *GPATCH8*. Indeed, single-gene validation experiments showed that loss of *PTEN*, *EPHA3*, *GPATCH8*, or *STK11* was individually sufficient to induce tumor formation in Barrett’s-derived epithelial cells, providing direct functional evidence for their tumor suppressive roles during EAC initiation.

While many of these genes have established or emerging roles in tumor biology in other cancer types, their specific contribution to EAC pathogenesis has remained poorly characterized. PTEN, a canonical tumor suppressor and negative regulator of PI3K/AKT signalling, is frequently altered across multiple cancers (Chalhoub and Baker 2009, Khasarah, Nabilsi et al. 2025). Similarly, together with our recent findings (Milne, Wu et al. 2026), *in vivo* evidence from this study showed that *PTEN* loss promotes tumorigenic transformation of premalignant Barrett’s epithelial cells through activation of PI3K/AKT/mTOR-associated programs.

The identification of EPHA3-associated signalling alterations is particularly interesting given the context-dependent roles of Eph receptors in cancer, where they can function as either tumor suppressors or oncogenes (Lv, Wang et al. 2018, Chen, Lu et al. 2019). Consistent with a tumor-suppressive role in Barrett’s tumorigenesis, EPHA3 loss strongly promoted tumor formation and was associated with coordinated transcriptomic and proteomic changes characterized by metabolic rewiring, including alterations in nucleic acid and nutrient metabolism pathways, together with suppression of epithelial integrity, and cell adhesion programs.

STK11 is a master regulator and “energy sensor” of the cell. Mechanistically, *STK11* loss impairs AMPK activation, leading to sustained mTORC1 signalling and enhanced macromolecule synthesis (Shaw, Bardeesy et al. 2004, Hardie 2005). This metabolic rewiring is further supported by HIF-1α-driven aerobic glycolysis (Faubert, Vincent et al. 2014). In our study, loss of STK11 induced EAC tumor formation and drove transcriptional and proteomic changes associated with multiple metabolic pathways, including cellular respiration, nucleic acid metabolism, and stress response pathways. However, STK11-deficient cells required the longest time to form tumours, suggesting that STK11 loss alone is insufficient to rapidly drive malignant transformation and likely depends on additional cooperating genetic alterations to accelerate tumor development.

A particularly intriguing finding from our screen was the identification of GPATCH8 as a potent tumor suppressor in this disease context. Although GPATCH8 is poorly characterized and has limited prior links to cancer, both *in vivo* screening and single-gene validation demonstrated that its loss initiate EAC tumor formation. Multi-omics profiling of GPATCH8-deficient cells revealed disruption of key programs including cell cycle regulation, DNA repair, and RNA metabolism, nuclear processes, cellular organization, and inflammatory responses. Among the proteins altered following GPATCH8 loss, the upregulation of EGFR, PTGS2, and MUC1 is particularly notable. Coordinated activation of EGFR and PTGS2 has been implicated in the progression of Barrett’s oesophagus to EAC, where PTGS2 promotes inflammation, angiogenesis, and resistance to apoptosis, while EGFR stimulates mitogenic signalling and cellular proliferation (Li, Wo et al. 2006, Wang, Xia et al. 2011, Korbut, Krukowska et al. 2022). Increased MUC1 expression may further potentiate this signalling network by stabilizing EGFR at the cell surface and enhancing downstream pathways involved in tumour cell migration and invasion (Butt, Pye et al. 2017, Du, Karatekin et al. 2025). Concomitantly, GPATCH8 deficiency was associated with reduced expression of ERRFI1, a well-established negative regulator of EGFR signalling, and its downregulation may permit sustained EGFR activation (Izumchenko and Sidransky 2015).

In addition to tumour initiation, our study addressed the clinically important question of intrinsic chemotherapy resistance. Resistance to platinum- and taxane-based chemotherapy remains a major barrier to effective treatment of EAC (Mao, Zeng et al. 2021), yet the underlying genetic determinants remain incompletely understood. Using pooled CRISPR screening under carboplatin and paclitaxel selection, we identified a set of shared resistance-associated tumour suppressor genes, including *NIPBL*, *PBRM1*, *AXIN1*, *RNF43*, *TGFBR2*, and *FAM196B*. Many of these genes regulate chromatin remodelling, WNT signalling, or TGF-β signalling pathways that have previously been implicated in therapeutic adaptation, cellular stress tolerance, and chemotherapy resistance (Cui, Jiang et al. 2012, Yuan, Tao et al. 2020, Lv and Xu 2021, Li, Huang et al. 2024). Among the shared resistance-associated genes, *NIPBL* emerged as a particularly compelling candidate, as its loss conferred resistance to both carboplatin and paclitaxel and was also associated with poorer overall survival in patients receiving platinum-based chemotherapy. *NIPBL* encodes a cohesin-loading factor involved in chromosome segregation, DNA replication, and DNA damage repair (Gao, Zhu et al. 2019), suggesting that disruption of cohesin-associated pathways may promote survival under chemotherapy-induced genotoxic stress. Importantly, although our functional screens were performed using carboplatin and paclitaxel, clinical validation was undertaken in patients treated with docetaxel, cisplatin, and 5-fluorouracil (DCF). Despite differences between these regimens, the overlap in drug class and mechanism supports the translational relevance of our findings. The association between low *NIPBL*, *TGFBR2*, and *RPL22* expression and poorer survival in DCF-treated patients therefore suggests that these tumour suppressors may contribute more broadly to resistance against platinum- and taxane-based therapies in EAC rather than resistance to a single agent alone.

In conclusion, this study provides a comprehensive functional framework linking recurrent genetic alterations in EAC to tumor initiation, transcriptional state remodelling, and chemotherapy resistance. We demonstrate that genetically diverse tumor suppressor perturbations converge onto a limited number of biologically coherent transcriptional programs. These shared transcriptional states provide a framework for future efforts to develop targeted therapies that can be applied across multiple patients exhibiting similar molecular phenotypes, irrespective of the specific underlying genetic alteration. Such an approach has the potential to overcome the extensive genomic heterogeneity of EAC by targeting common downstream programs that emerge from distinct tumor suppressor perturbations. Our findings expand the current understanding of EAC pathogenesis and resistance to chemotherapies, identify several previously underappreciated candidate tumor suppressors, and provide a resource for future mechanistic and translational investigations aimed at improving early detection and therapeutic intervention in EAC.

## METHODS

### Animal studies

All animal experiments were performed in accordance with the National Health and Medical Research Council Australian Code of Practice for the Care and Use of Animals for Scientific Purposes, with approval from the Peter MacCallum Cancer Centre Animal Ethics Committee (#E698). Female NOD.Cg-Prkdcscid Il2rgtm1Wjl/SzJ (NSG) mice, aged 4-6 weeks, were socially housed in individually ventilated cages within temperature and light cycle controlled, specific pathogen-free rooms at the Peter MacCallum Cancer Centre. Mice had ad libitum access to food, water, and nesting materials and were monitored at least weekly for general wellbeing.

### Cell culture and viability assay

The Barrett’s esophagus epithelial cell lines CP-B (CP-52731; RRID: CVCL_C452) and CP-D (CP-18821; RRID: CVCL_C454) were provided by Professor Peter Rabinovitch (University of Washington, Seattle, WA). These lines were originally established from regions of Barrett’s esophagus with high-grade dysplasia (Palanca-Wessels, Barrett et al. 1998), and represent relevant models for investigating Barrett’s tumorigenesis. Both CP-B and CP-D cell lines harbor mutations in TP53 and CDKN2A and exhibit genomic copy number alterations (Palanca-Wessels, Barrett et al. 1998, Suchorolski, Paulson et al. 2013). Cells were cultured in MCDB-153 medium (Sigma-Aldrich) supplemented with 400 ng/mL hydrocortisone, 20 mg/mL adenine, 1x insulin-transferrin-sodium selenite supplement, 10 nmol/L cholera toxin, 375 ng/mL fluconazole, penicillin/streptomycin (100 U/mL and 100 μg/mL, respectively) (all from Sigma-Aldrich), 20 ng/mL epidermal growth factor (PeproTech), 84 mg/mL bovine pituitary extract (Banksia Scientific), 5% foetal bovine serum (FBS, Gibco), and 4mM L-glutamine (Gibco). HEK293T/17 cells (CRL-11268G-1; RRID: CVCL_UE07) were obtained from the American Type Culture Collection (ATCC, USA) and maintained in DMEM high-glucose medium (Sigma-Aldrich) supplemented with 10% FBS (Gibco) and 2 mM L-glutamine (Gibco). All cell lines were maintained at 37 °C in a humidified incubator with 5% CO_2_. Cell line identity was verified by short tandem repeat (STR) profiling conducted by the Australian Genome Research Facility (AGRF) using the Promega GenePrint 10 system. All cultures were routinely tested for mycoplasma contamination and confirmed negative by PCR (Genotyping Core Facility, Peter MacCallum Cancer Centre, Australia). For cell viability assays, cells were seeded in 150 mm tissue culture dishes (Corning) at 3.5 × 10⁶ cells per dish to mimic the culture conditions used in the chemotherapy resistance screen, with three dishes per condition. After 96 hours of treatment with vehicle (normal saline), 35 μM carboplatin (Accord Healthcare, Australia; Cat. No. 215854), or 3 μM paclitaxel (Sandoz, Australia; Cat. No. 98551), cells were trypsinised, stained with trypan blue (Gibco), and live and dead cells were counted using a Countess cell counter (Thermo Fisher).

### Lentiviral production and infection

HEK293T/17 cells were transfected overnight at 371°C with 101µg of the vector plasmid, 3 µg of pMD2.G (Addgene #12259), and 6 µg of psPAX2 (Addgene #12260) packaging plasmids using polyethylenimine (PEI; 11mg/mL stock) at a 3:1 w/w ratio relative to DNA, as described (Iggo and Richard 2015). Following overnight incubation, the medium was replaced with Opti-MEM (Gibco) supplemented with 20 mM HEPES. Conditioned medium containing viral particles was collected 481hours post-transfection, filtered through a 0.45 µm membrane, centrifuged at 2,1001×1g, and concentrated ∼200-fold using Vivaspin20 columns (GE Healthcare) according to the manufacturer’s instructions. Target cells were transduced with the concentrated lentivirus in the presence of 8 µg/mL polybrene at a multiplicity of infection (MOI) of 1-2 infectious units per cell, except for pooled CRISPR library experiments where MOI was < 0.3. Transduced cells were subsequently selected with the appropriate antibiotic, added to culture medium at standard concentrations and durations as required for each plasmid.

### Generation of Cas9-expressing CPB and CPD cell lines

CPB and CPD cells were transduced with lentiviral particles containing FUCas9Cherry plasmid (Addgene #70182) as described above. Following transduction, cells were harvested, resuspended in medium containing 1% FBS and 0.25 mM EDTA to minimize clumping, and filtered through 40 µm cell strainers to remove aggregates. Single-cell clones were then isolated by fluorescence-activated cell sorting (FACS) using a BD FACSAria Fusion-5 Cell Analyzer based on mCherry expression and deposited into 96-well plates containing complete MCDB-153 medium. Individual clones were expanded under standard culture conditions and assessed for Cas9 nuclease activity using an EGFP reporter assay (Addgene #59702), in which EGFP expression is lost upon successful Cas9-mediated cleavage. Clones were transduced with the EGFP reporter and subjected to puromycin selection (2 µg/mL, Gibco) for seven days, with media refreshed every 3 days to remove non-viable cells. EGFP fluorescence was assessed using a BD FACSCanto II flow cytometer. A single clone from CPB (designated Cas9-CPB) and CP-D (designated Cas9-CPD) showing minimal or absent EGFP signal, corresponding to > 95% Cas9 activity, was selected for further expansion and subsequent experiments.

### Pooled CRISPR-Cas9 library design and transduction

Single guide RNAs (sgRNAs) targeting a panel of fifty-six putative EAC tumour suppressor genes (GSE335866) were designed in collaboration with Skyang Bio (formerly Transomic Technologies, Alabama, USA) using the Croatan sgRNA design algorithm and cloned into the pCLIP-Dual-SFFV-ZsGreen-Puromycin (V163), as described previously (Erard, Knott et al. 2017). Each gene was targeted with six sgRNAs incorporated as pairs within dual-guide vectors (three clones per gene), combining higher- and lower-scoring guides to balance predicted on-target efficiency and maximize knockout performance. Cas9-CPB and Cas9-CPD cells were transduced with the pooled CRISPR knockout library, containing in total 178 sgRNAs targeting the 56 genes and 10 non-targeting control sgRNAs (scrambled, SCR), at a MOI < 0.3 to ensure predominantly single-vector integration per cell. Twenty-four hours post-transduction, the medium was replaced with medium containing puromycin (2 µg/mL; Gibco), and selection was carried out for seven days. Following selection, cells were trypsinised, with at least 20,000 cells collected for single-cell sequencing, while the remaining population was expanded for subsequent *in vivo* tumorigenesis or *in vitro* drug screen experiments.

### Pooled CRISPR-Cas9 loss-of-function *in vivo* screen

On the day of injection, 5 × 10⁶ cells from the pooled CRISPR library transduced Cas9-CPB and Cas9-CPD cells were collected to establish the T0 baseline. An equivalent number of cells were suspended in Matrigel (1:1 ratio, Corning) and injected subcutaneously into the flank region of NSG mice (4 per cell line). Mice were monitored weekly for tumor formation, and tumor size were measured using digital callipers. Tumours were collected once they reached a volume of ∼500 mm³, a size chosen to provide sufficient material for DNA extraction and sequencing while minimizing clonal dominance. Small portions of each tumor were fixed in neutral-buffered formalin and processed for immunohistochemistry, whereas the remaining tissue was snap-frozen for DNA extraction.

### *In vitro* CRISPR screening with chemotherapy treatment

Cas9-CPB and Cas9-CPD cells transduced with the pooled CRISPR library were seeded in 150 mm tissue culture dishes (Corning) at 3.5 × 10⁶ cells per dish, with three dishes per condition and three technical replicates. After 96 hours of treatment with vehicle (normal saline), 35 μM carboplatin (Accord Healthcare, Australia; Cat. No. 215854), or 3 μM paclitaxel (Sandoz, Australia; Cat. No. 98551), cells from the three dishes of each condition within each replicate were pooled. Genomic DNA was then extracted as described below.

### Genomic DNA processing and high-throughput sequencing

Genomic DNA was extracted from snap-frozen cell pellets or tumor samples using the DNeasy Blood and Tissue Kit (Qiagen). Briefly, snap-frozen cell pellets were thawed on ice and resuspended in 200 μL 1x PBS with 20 μL Proteinase K. Tumours were cut into pieces (∼25 mg) and homogenized using PowerBead Ceramic Tubes and a MO Bio PowerLyzer 24 homogenizer in 180 μL tissue lysis buffer. Following homogenization, 20 μL Proteinase K was added, and samples were incubated at 56°C until complete lysis. The homogenates were centrifuged at 10,000 × g for 20 min, and the supernatants collected and processed according to the manufacturer’s instructions. Next-generation sequencing libraries were generated using a two-step PCR approach. PCR1 was performed with Herculase II Fusion DNA Polymerase (Agilent). The reaction mix contained 10 μL 5× Herculase reaction buffer, 0.5 μL 25 mM dNTP mix, 1.25 μL forward primer (pCLIP-Dual_vector_F), 1.25 μL reverse primer (pCLIP-Dual_vector_R), 0.5 μL Herculase II Fusion polymerase, 250 ng DNA, and PCR-grade water to a final volume of 50 μL. PCR reaction was run under the following conditions: initial denaturation at 98°C for 15 min; 25 cycles of 95°C for 15 sec, 57°C for 45 sec, and 72°C for 45 sec; followed by a final extension at 65°C for 5 min. PCR products were separated on a 2% agarose gel, extracted, and purified using the NucleoSpin® Gel and PCR Clean-up Kit (Macherey-Nagel) according to the manufacturer’s instructions. PCR1 reactions from each sample were pooled and stored at −20°C. PCR2 was performed with KOD Hot Start Polymerase (Novagen). The reaction mix contained 5 μL 10× KOD reaction buffer, 5 μL 25 mM dNTP mix, 2 μL 10 μM of forward (P5) and reverse (P7) indices and sequencing adapters, 2.5 μL dimethyl sulfoxide (DMSO), 4 μL 25 mM MgSO₄, 1.5 μL KOD polymerase, 750 ng PCR1 amplicon, and PCR-grade water to a final volume of 50 μL. PCR2 was run under the following cycling conditions: initial denaturation at 95°C for 2 min; 6 cycles of 95°C for 20 sec, 65°C for 10 sec, and 70°C for 15 sec; 16 cycles of 95°C for 20 sec and 70°C for 25 sec; and a final extension at 70°C for 2 min. PCR2 products were purified using the NucleoSpin® Gel and PCR Clean-up Kit (Macherey-Nagel) according to the manufacturer’s instructions. DNA libraries were sequenced on the Illumina NovaSeq X Sequencing System at the AGRF. All primer sequences are provided in *Supplementary Table 1*.

### CRISPR screen data analysis

Sequencing reads were aligned to the reference gRNAs library using the Galaxy bioinformatic analysis platform (Afgan, Baker et al. 2018). Guide counts for each gene were obtained using MAGeCK count (Galaxy Version 0.5.9.2.4). To identify gene knockouts that conferred increased resistance to chemotherapy, MAGeCK test (Galaxy Version 0.5.9.2.1) was applied to compare treated versus control samples, highlighting positively selected knockouts. For *in vivo* tumor samples, enriched sgRNAs were determined by comparing tumours collected at endpoint to the corresponding T0 samples obtained on the day of injection. All analyses were performed using default parameters unless otherwise specified. Data visualization and graphical representation were performed in RStudio (version 2025.05.1) using the ggplot2, dplyr, and ggrepel packages.

### Generation of CRISPR individual gene knockout cells

Individual knockout Cas9-CPB cell lines for *GPATCH8*, *STK11*, and *EPHA3* were generated using the pCLIP-Dual-SFFV-ZsGreen-Puromycin sgRNA expression vector, with cells transduced with at least two independent sgRNAs (GPATCH8: TEDH-1032437 and TEDH-1032441; STK11: TEDH-1077640 and TEDH-1077641; EPHA3: TEDH-1025049 and TEDH-1025050, *Supplementary Table 1*). For the *PTEN* knockout cell line, Cas9-CPB cells were transduced with a PTEN-targeting sgRNA cloned into the lentipuro-guide expression vector (Addgene # 52963), which was generated in our laboratory as described earlier (Milne, Wu et al. 2026). . For western blot, and *in vivo* experiments, EPHA3, GPATCH8, and STK11 samples were pooled from two independent sgRNAs.

### Western blotting

Cell lines were seeded in 80 mm tissue culture dishes (Corning) at ∼60% confluency and allowed to attach for 48 hours before lysis in RIPA buffer (50 mM Tris-HCl pH 8.0, 150 mM NaCl, 1% NP-40, 0.5% sodium deoxycholate, 0.1% sodium dodecyl sulphate (SDS)) supplemented with phosphatase (PhosphoSTOP, Roche) and protease (cOmplete, Roche) inhibitors. Protein concentrations were determined using the Pierce™ BCA Protein Assay Kit (Thermo Fisher Scientific). Lysates (40 µg protein per sample) were denatured in 1× SDS sample buffer (313 mM Tris-HCl pH 6.8, 50% glycerol, 10% β-mercaptoethanol, 10% SDS, 0.05% bromophenol blue) at 95 °C for 5 min and resolved on 10% Bis-Tris SDS-PAGE gels. Proteins were transferred to membranes using standard wet transfer protocols, blocked for 1 hour in 5% skim milk in Tris-buffered saline (TBS) containing 0.1% Tween-20, and incubated overnight at 4°C with primary antibodies: anti-STK11 (Cell Signaling Technology, Cat# 27D1, 1:1000), anti-PTEN (Cell Signaling Technology, Cat# 9559, 1:1000), anti-EPHA3 (Santa Cruz Biotechnology, SC-514209, 1:1000), anti-GPATCH8 (Bioss, BS-13502R, 1:1000), anti-β-Actin (Sigma, A2228, 1:5000), and anti-Vinculin (Bio-Rad, MCA465GA, 1:10000) . HRP-conjugated anti-mouse (Bio-Rad, #1706516, 1:5000) or anti-rabbit (Bio-Rad, #1706515, 1:5000) secondary antibodies were used as appropriate, and signals were detected with Clarity Western ECL substrate (Bio-Rad) on a ChemiDoc MP imaging system (Bio-Rad).

### Histology and immunohistochemistry staining

Formalin-fixed, paraffin-embedded (FFPE) tissue blocks were sectioned at 4 µm using an RM2235 manual rotary microtome (Leica), mounted onto adhesive microscope slides (TRAJAN), and baked for 60 min at 60 °C before deparaffinisation in xylene and dehydration with 100% ethanol. Sections were stained with Lillie-Mayer’s haematoxylin (Australian Biostain), differentiated with 0.3% acid alcohol, and counterstained with 1% alcoholic eosin/phloxine (Australian Biostain). For immunohistochemistry, antigen retrieval was performed in citrate buffer (pH 6), followed by blocking with 3% hydrogen peroxide and 10% bovine serum albumin (BSA) in TBS containing 0.05% Tween-20 (TBS-T). Slides were incubated overnight with primary antibodies against human mitochondrial protein (Merck Millipore, MAB1273, 1:1000). The following day, sections were washed with TBS-T and incubated for 1 hour with DAKO Mouse Envision secondary antibody. Protein localisation was visualised using liquid 3,3’-diaminobenzidine (DAB) substrate chromogen mixture (Dako). Slides were counterstained with haematoxylin and scanned using the SLIDEVIEW VS200 scanner (Olympus) and visualized with Fiji software.

### RNA-sequencing and data analysis

Cells were seeded in 6-well tissue culture plates (Corning) at 60% confluency and allowed to attach for 48 hours prior to RNA extraction using the NucleoSpin RNA kit (Macherey-Nagel, Germany) following the manufacturer’s instructions. Three biological replicates were collected for RNA sequencing representing independent consecutive passages. RNA concentration and purity were measured with a NanoDrop1000 spectrophotometer (Thermo Fisher Scientific). For each sample, 10 µg of RNA was submitted to AGRF for library preparation and sequencing. Libraries were prepared using the TruSeq Stranded mRNA Library Prep kit (Illumina) and sequenced on an Illumina NovaSeq X platform with 150 bp paired-end reads. Sequencing quality was assessed using FastQC (Galaxy Version 0.74+galaxy1). Reads were aligned to the GRCh38 reference genome (Ensembl release 101) with HISAT2 (Galaxy Version 2.2.1+galaxy1), and gene features were quantified with featureCounts (Galaxy Version 2.0.3+galaxy2). Differential expression analysis was performed using edgeR (Galaxy Version 3.36.0+galaxy4), and pathway enrichment analysis was conducted in RStudio (version 2025.05.1) using the ClusterProfiler (Yu, Wang et al. 2012), and results were visualized with ggplot2.

### Perturb-seq experiments, library preparation, and sequencing

Cells from the pooled CRISPR-Cas9 library transduction were used for downstream single-cell RNA sequencing. Seven days post-transduction, cells were resuspended in 1× PBS with 0.04% BSA according to the 10x Genomics Single Cell Protocols Cell Preparation Guide (CG00053 Rev C). Cells were encapsulated into droplet emulsions using the Chromium Controller (10x Genomics) with Chromium Single-Cell 31 Gel Beads v3 (PN2000164), aiming to recover ∼10,000 cells per cell line, with an average of ∼165 cells per gene. Library preparation was performed using the 10x Genomics Chromium Single Cell 31 Reagent Kits v3 with Feature Barcode technology for CRISPR screening (CG000184 Rev C), with magnetic selections conducted on an Alpaqua Catalyst 96 plate. mRNA and sgRNA libraries were sequenced on a NovaSeq 6000 (Illumina) following the 10x Genomics User Guide. Raw sequencing data were processed using the Cell Ranger v6.0.2 pipeline, which uses the STAR aligner to map reads to the GRCh38-3.0.0 human reference genome and to the CRISPR guide library. Cells were filtered to remove low-quality cells (<10,000 UMI counts) and those with high mitochondrial content (> 5%). Only cells harbouring a single sgRNA were retained for downstream analysis. Downstream analysis was conducted in RStudio (version 2025.05.1). The gene expression and guide count matrices were processed using Seurat for normalization and scaling. Pseudobulk differential expression analysis between perturbed cells and scrambled controls was performed using Seurat’s FindMarkers, with significance defined as p-value < 0.05. Pathway enrichment analysis was conducted using the ClusterProfiler (Yu, Wang et al. 2012) or Metascape (Zhou, Zhou et al. 2019). For each perturbation, enrichment results from the two cell lines were combined: if a pathway was regulated in the same direction in both cell lines, the FDR-values were averaged; if it was regulated in only one cell line, the original FDR-value was retained. Gene functional similarity was quantified using a signed Jaccard similarity score, representing the proportion of pathways regulated in the same direction between gene pairs. Heatmaps and network graph were generated using ComplexHeatmap, igraph, tidygraph, and ggraph packages.

### Mass spectrometry sample preparation and analysis

Proteomic samples were prepared using an S-Trap micro spin column digestion workflow (HaileMariam, Eguez et al. 2018). Briefly, 1 × 10⁶ Cas9-CPB cells per condition were lysed in SDS/TEAB buffer supplemented with benzonase and protease inhibitors. Protein concentration was determined Pierce™ BCA Protein Assay Kit (Thermo Fisher Scientific), and 100 µg of protein per sample was subjected to reduction (TCEP) and alkylation (iodoacetamide) prior to acidification and protein capture on S-Trap columns. Proteins were digested overnight with trypsin (1:20 w/w) at 37°C, and peptides were sequentially eluted using TEAB, formic acid, and acetonitrile/formic acid buffers. Lyophilized peptides were resuspended in 2% acetonitrile/0.05% trifluoroacetic acid (TFA), centrifuged at 14,000 × g for 10 min, and subjected to LC–MS/MS analysis. Mass spectrometry was performed at the Bio21 Institute (University of Melbourne) using a data-independent acquisition (DIA) platform. Raw data were processed and quantified using Analyst Suite (Monash Proteomics Platform).

### Public Datasets

The Cancer Genome Atlas (TCGA) Firehose Legacy dataset was accessed through cBioPortal (Cerami, Gao et al. 2012, Gao, Aksoy et al. 2013). The EAC ICGC dataset, including mutation, copy number alteration, and clinical data, was provided by Professor Nicola Waddell (QIMR Berghofer Medical Research Institute, Brisbane, Australia). For the chemotherapy resistance analysis, patient data were primarily obtained from the DOCTOR clinical trial (ANZCTR - ACTRN12609000665235). EAC patients enrolled in the DOCTOR clinical trial, as previously described (M. Naeini, Newell et al. 2023), received pre-operative chemotherapy and/or radiotherapy. Processed mRNA expression and clinical annotation data were provided by Professor Nicola Waddell (QIMR Berghofer Medical Research Institute, Brisbane, Australia).

### Statistical analysis

Statistical analysis was performed in RStudio (version 2025.05.1) and GraphPad Prism 11.0.1. Detailed statistical methods are described in the figure legends or specific methods. Patient survival differences were estimated using Kaplan-Meier method and significance was examined using the log-rank test.

---

**Supplementary Table 1. Sequences of primers and sgRNAs used in this study.**

**Supplementary Table 2. Pathway enrichment analysis of perturbation-defined clusters.** The table summarizes significantly enriched pathways associated with each cluster, including pathway name, corresponding gene perturbation, leading-edge genes contributing to pathway enrichment, and the direction of pathway regulation (upregulated or downregulated).

**Supplementary Table 3. Protein profiling of Cas9-CP-B cells following knockout of *EPHA3, GPATCH8,* and *STK11*.**

## Acknowledgements

The authors acknowledge the Molecular Genomics Core (RRID:SCR_025695), the Centre for Advanced Histology and Microscopy (RRID:SCR_025432), the Victorian Centre for Functional Genomics (RRID:SCR_025582), the Research Computing, and the Cancer Research Animal Core Facilities at Peter MacCallum Cancer Centre for access and support. We also acknowledge the technical and bioinformatics support provided by Brodie Bilston, Dr Sandun Rajapaksa, and Dr Hao Nathan Feng.

---

**Supplementary Figure 1.**
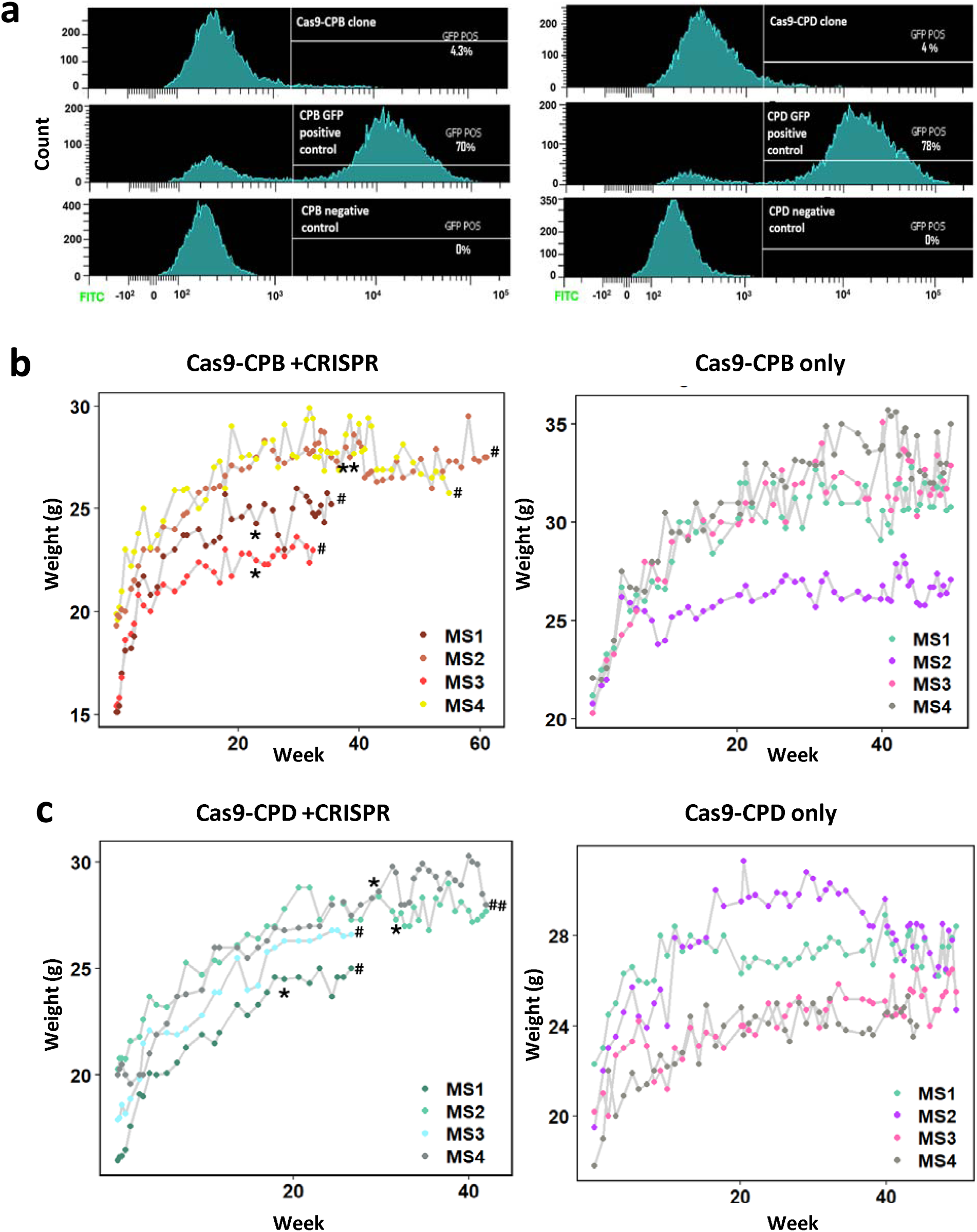
Cas9 function validation and body weight monitoring in mice from in vivo screen. **(a)** Flow cytometry analysis of EGFP reporter cleavage in ectopic mCherry-Cas9 expressing CPB and CPD single-cell clones (Cas9-CPB, Cas9-CPD). Efficient Cas9 activity was demonstrated by less than 5% GFP-positive cells remaining (top plots). Positive and negative controls confirmed transduction efficiency and gating accuracy (middle and bottom plots, respectively). **(b)** Body weight monitoring of mice (MS) injected with CRISPR library transduced Cas9-CPB cells or Cas9-CPB cells alone. *: Week of tumor appearance post-injection. #: Week of culling for tumor analysis. Mice injected with Cas9-CPB cells alone did not develop tumours and were culled after one year in accordance with project ethical guidelines. **(c)** Body weight monitoring of mice (MS) injected with Cas9-CPD cells, presented as in (b).

**Supplementary Figure 2.**
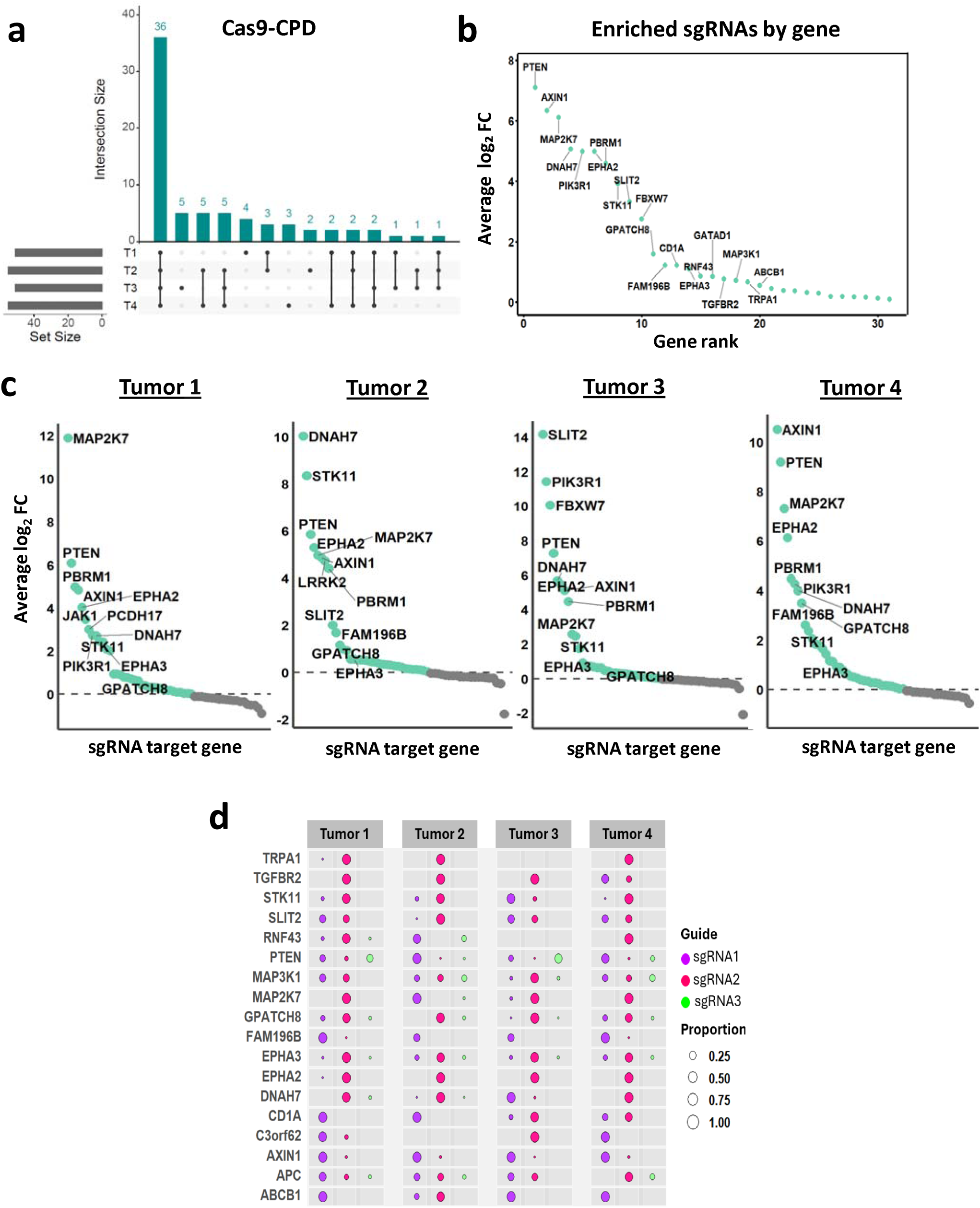
Identification of tumor suppressors whose loss drives BE cell transformation. **(a)** UpSet plot showing the number of enriched sgRNAs shared across four Cas9-CPD tumours. **(b)** Average log_2_ FC of tumor-enriched sgRNAs in Cas9-CPD tumours, grouped by gene and relative to baseline (T0) cells. Displayed genes were identified in tumours from at least three mice. **(c)** MAGeCK plots showing log_2_ FC of enriched sgRNAs grouped by gene in four independent Cas9-CPD tumours. Top-enriched genes are labelled. **(d)** Enrichment of independent sgRNA clones across four tumours, with each gene targeted by three pairs of sgRNA. Dot size reflects the proportion of total reads per gene.

**Supplementary Figure 3.**
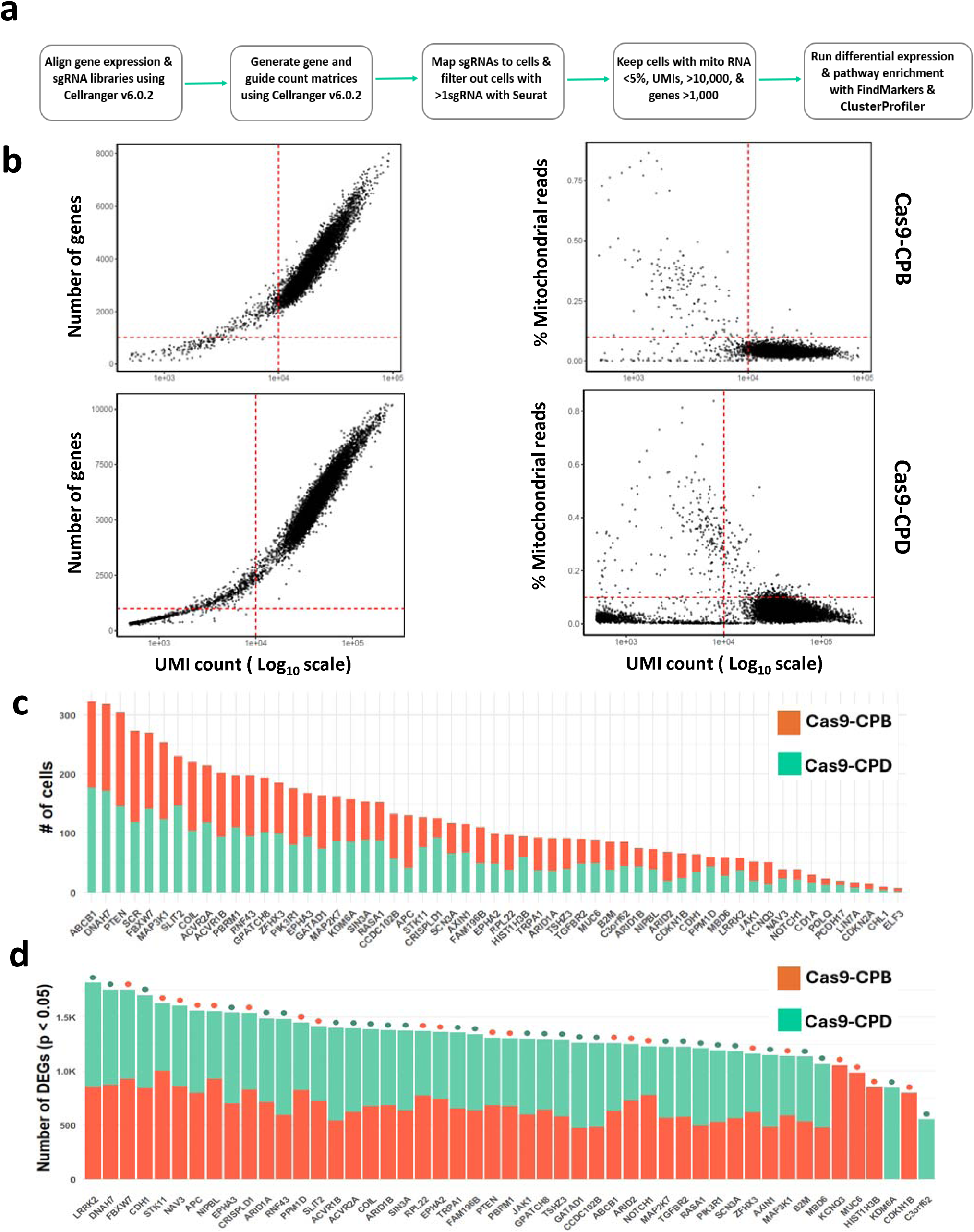
Data analysis and quality filtering of Perturb-seq results shown in Figure 4. **(a)** Schematic overview of data alignment, cell calling, sgRNA assignment, and filtering. **(b)** Dot plots show total UMI counts (log₁₀) versus gene number (cutoffs: 10,000 UMIs, 1,000 genes) and mitochondrial gene percentage (cutoff: <5%). **(c)** Stacked bar plot showing the number of cells per gene perturbation. **(d)** Stacked bars show differentially expressed genes (DEGs) counts per perturbation. Single-color bars mean data missing from one cell line due to low cells or poor knockdown of target gene. Dots above indicate the cell line with the higher number of DEGs.

**Supplementary Figure 4.**
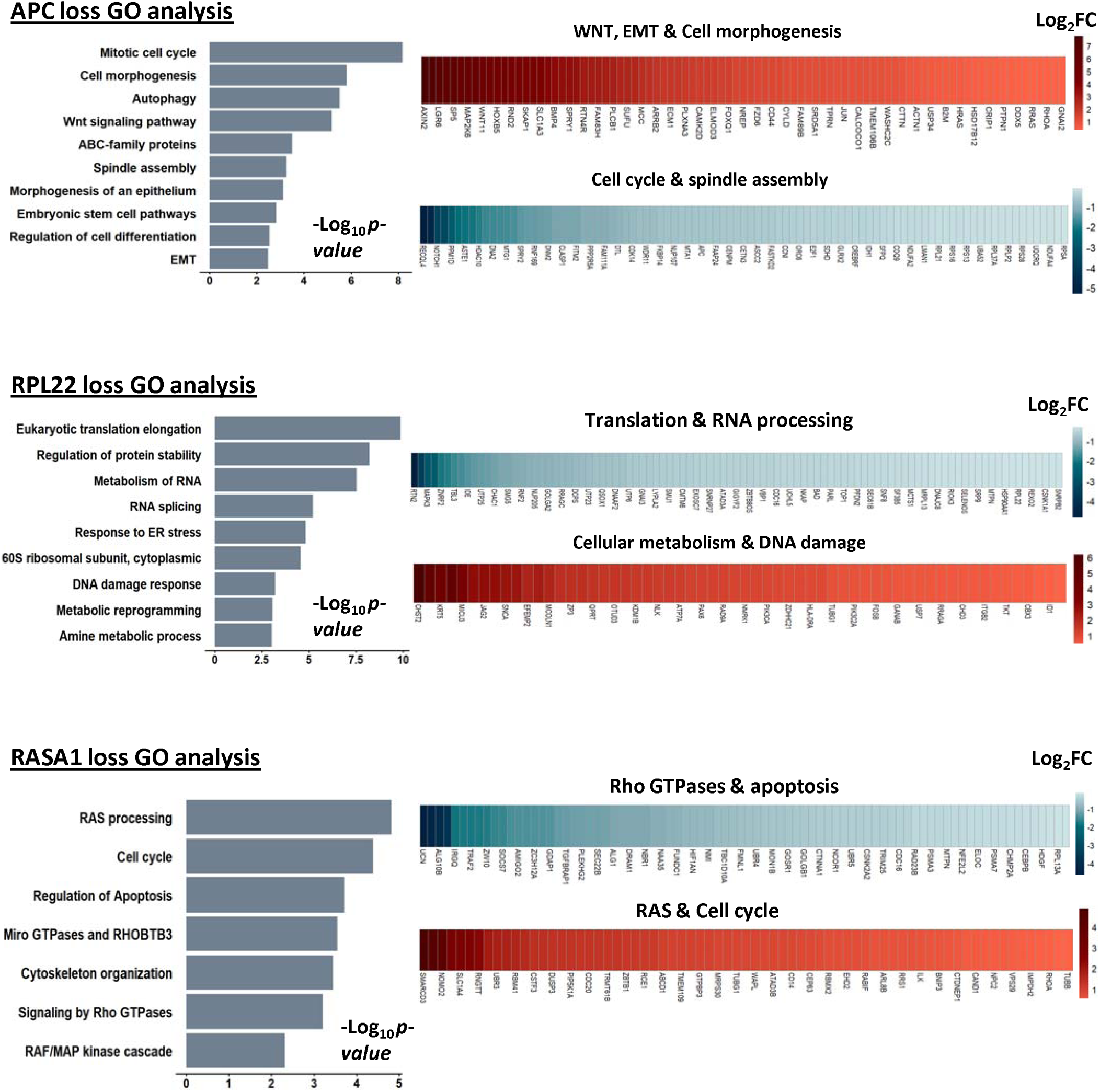
CRISPR-mediated knockout of well-characterised genes produces expected transcriptomic signatures in Perturb-seq. Bar plots show pathway enrichment analysis following knockout of APC, RPL22, and RASA1, alongside heatmaps showing representative examples of genes enriched in these pathways. Data from both CP-B and CP-D cell lines were combined.

**Supplementary Figure 5.**
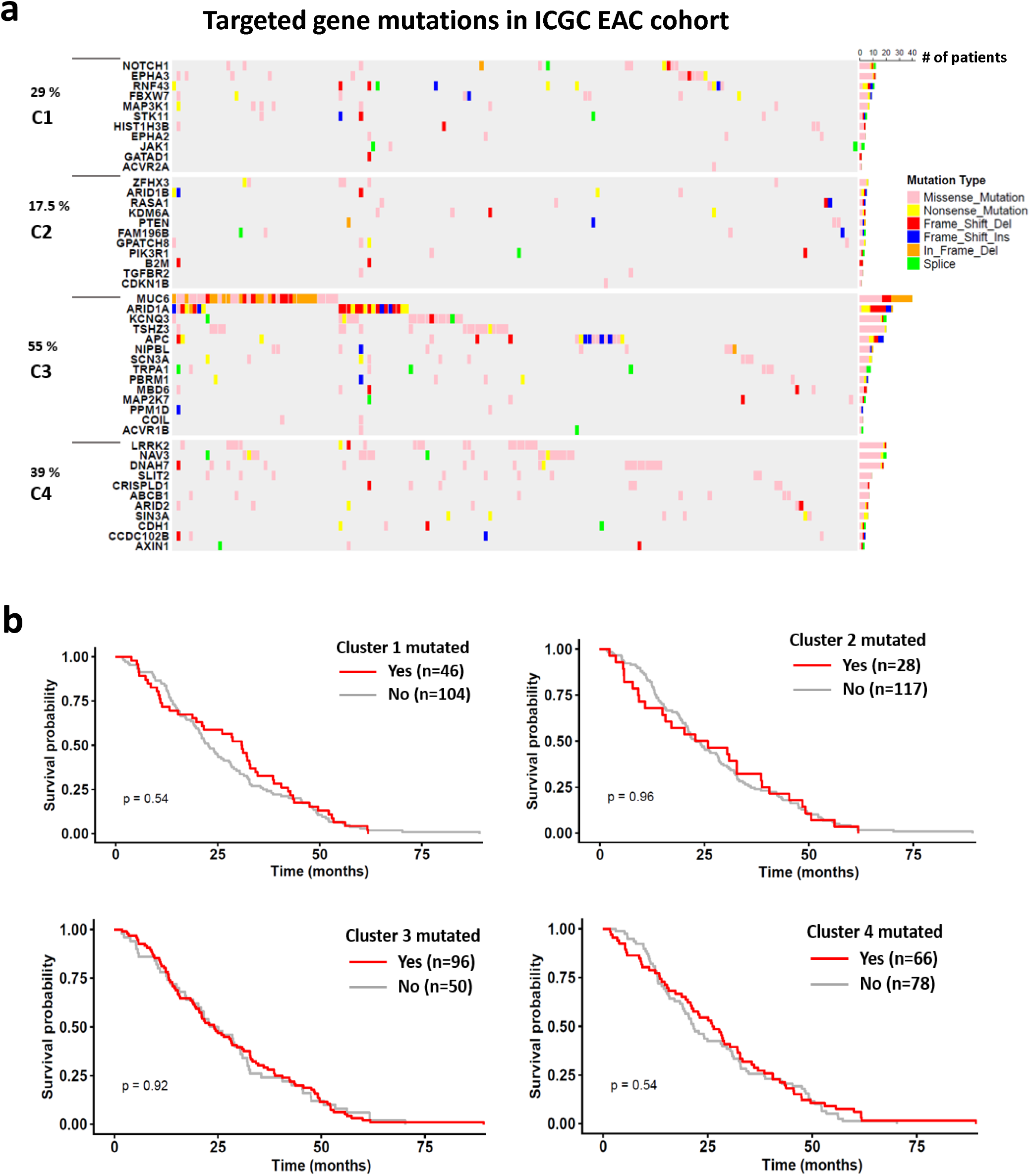
Representation of major transcriptional clusters within the ICGC EAC clinical cohort. **(a)** Oncoprint heatmap showing the percentage of EAC patients (indicated in black) from the ICGC cohort harbouring genetic alterations enriched in clusters C1-C4. The plot displays mutation types, the number of patients per alteration, and overlapping mutations across samples. **(b)** Kaplan-Meier survival curves of ICGC EAC patients stratified by cluster status. The red curve (“Yes”) represents patients with a tumor harbouring alteration in one or more genes within the assigned cluster, including missense, nonsense, frameshift, splice-site mutations, and deep deletions, whereas the grey curve (“No”) represents patients with a tumor without alterations in any gene within the assigned cluster.

**Supplementary Figure 6.**
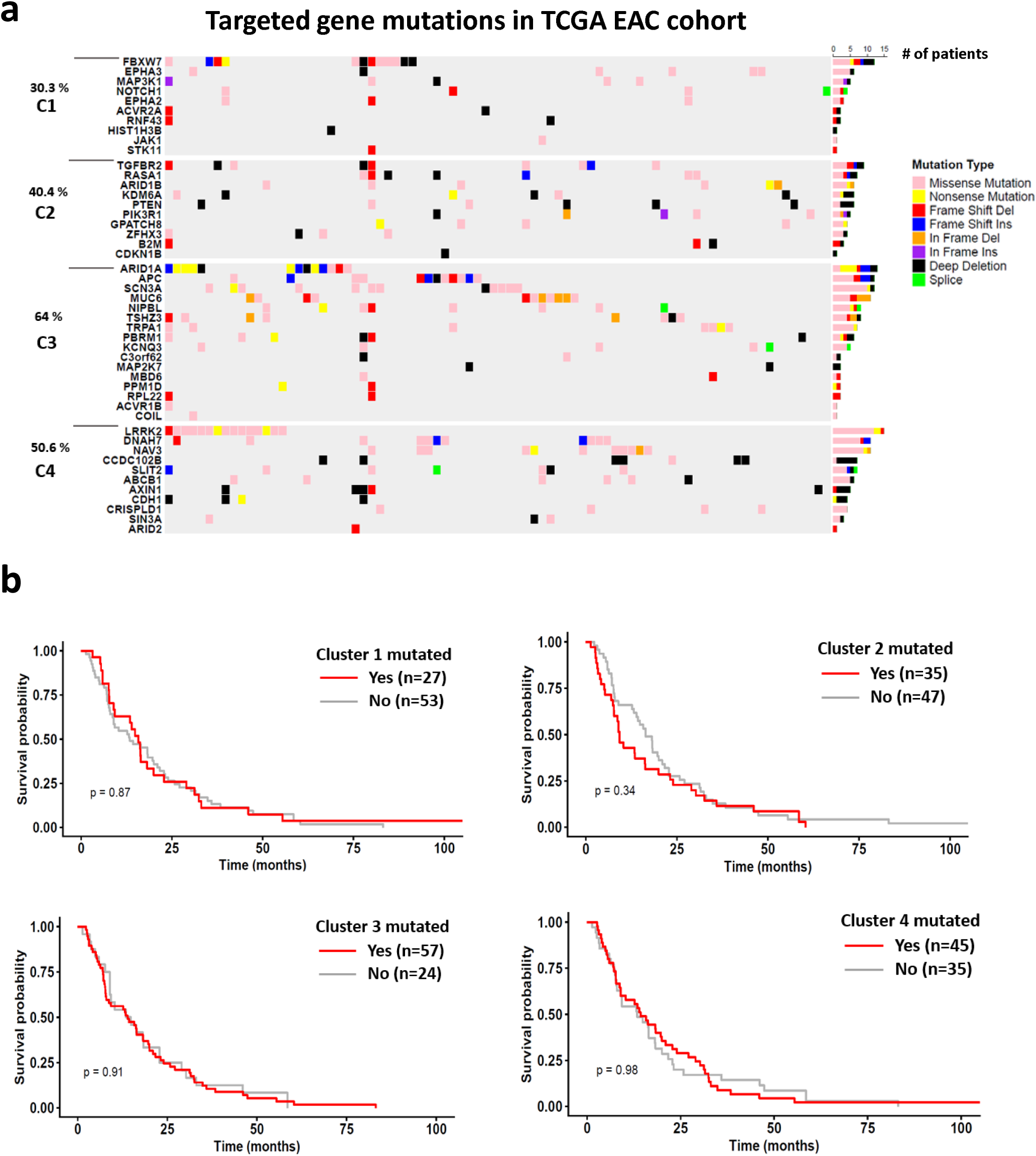
Representation of major transcriptional clusters within the TCGA EAC clinical cohort. **(a)** Oncoprint heatmap showing the percentage of EAC patients (indicated in black) from the TCGA cohort harbouring genetic alterations enriched in clusters C1-C4. The plot displays mutation types, the number of patients per alteration, and overlapping mutations across samples. **(b)** Kaplan-Meier survival curves of TCGA EAC patients stratified by cluster status. The red curve (“Yes”) represents patients with a tumor harbouring alteration in one or more genes within the assigned cluster, including missense, nonsense, frameshift, splice-site mutations, and deep deletions, whereas the grey curve (“No”) represents patients with a tumor without alterations in any gene within the assigned cluster.

**Supplementary Figure 7.**
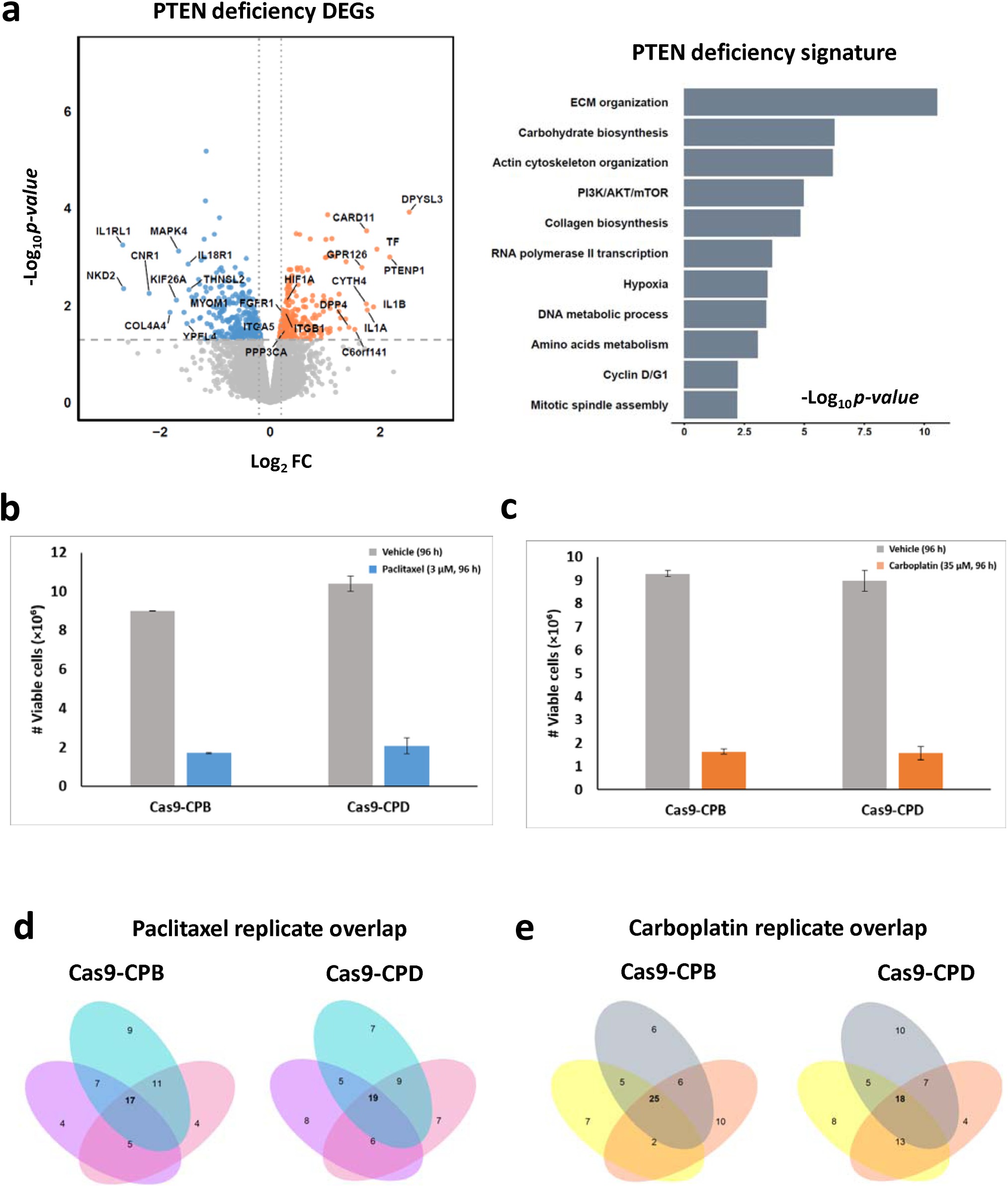
PTEN loss bulk RNA-seq analysis and chemotherapy resistance screen. **(a)** Bar and volcano plots showing pathway enrichment analysis and differentially expressed genes (from bulk RNA-seq) in Cas9-CPB cells following PTEN loss. **(b-c)** Bar charts showing the number of viable cells following treatment with either vehicle, Paclitaxel (3 μM), or Carboplatin (35 μM) for 96 hours. Error bars represent the standard error of the mean (SEM) from three biological replicates. **(d-e)** Venn diagrams showing the number of sgRNAs shared among the three replicates of the Paclitaxel and Carboplatin screens in each cell line.

